# SnapFISH: a computational pipeline to identify chromatin loops from multiplexed DNA FISH data

**DOI:** 10.1101/2022.12.16.520793

**Authors:** Lindsay Lee, Hongyu Yu, Bojing Blair Jia, Adam Jussila, Chenxu Zhu, Jiawen Chen, Liangqi Xie, Antonina Hafner, Caterina Strambio-De-Castillia, Alistair Boettiger, Bing Ren, Yun Li, Ming Hu

## Abstract

Multiplexed DNA fluorescence in situ hybridization (FISH) imaging technologies have been developed to map the folding of chromatin fibers at tens of nanometer and tens of kilobase resolution in single cells. However, computational methods to reliably identify chromatin loops from such imaging datasets are still lacking. Here we present a Single-Nucleus Analysis Pipeline for multiplexed DNA FISH (SnapFISH), to process the multiplexed DNA FISH data and identify chromatin loops. SnapFISH can identify known chromatin loops from mouse embryonic stem cells with high sensitivity and accuracy. In addition, SnapFISH obtained comparable results of chromatin loops across datasets generated from diverse imaging technologies.

## 1. Introduction

How chromatin folds inside the nucleus is a fundamental question in the study of genome structure and function^1^. Disruption of chromatin organization can lead to gene dysregulation, and has been associated with a variety of human developmental disorders, neuropsychiatric diseases and cancers^2^. Different from proximity-ligation assays^3,4^ that infer the 3D genome through *indirect* measurement of DNA sequence contacts, chromatin tracing, as an emerging microscopy-based technology, can visualize bright spots corresponding to individual targeted genomic segments arrayed along chromatin fibers, and map their physical location in threedimensional space. This rich imaging data permits the *direct* measurement of Euclidean distances between targeted genomic segments of interest - such as promoters and distal *cis*-regulatory elements - allowing an intimate look into the organization of chromosomes^5^. During the last decade, a number of chromatin tracing technologies have emerged, including multiplexed DNA fluorescence in situ hybridization (FISH)^6^, DNA-MERFISH^7^, DNA seqFISH+^8,9^, ORCA^10^, MINA^11^, Hi-M^12^, OligoFISSEQ^13^ and IGS^14^ (more details can be found in two recent review articles^5,15^). These techniques can resolve the spatial location of discrete targeted genomic segments with tens of nanometer precision in single cells. They have been used to image the entire mammalian genome at megabase resolution^8,9^, one full chromosome at 50 kilobase (Kb) resolution^7^, and a few selected mega-base regions at 5Kb ~ 25Kb resolution^8–10,16^, promising to uncover novel insights into chromatin folding and its role in gene regulation^5^.

Chromatin loops are a key structural feature of chromatin spatial organization, and may serve as the structural basis of gene regulation. Originally discovered from bulk Hi-C data^4,17^ as “dots” at the corners of topologically associating domains, and recently identified from single cell Hi-C data^18,19^ and imaging data^16^, chromatin loops are defined as pairs of genomic loci with closer spatial proximity compared to other pairs of loci in the local neighborhood region^4,17^. Chromatin loops between enhancers and promoters have been used to infer target genes for distal enhancers^20^. Chromatin loops between CTCF binding sites lead to the formation of topological associating domains^21^, which are megabase (Mb) sized chromatin domains constraining enhancer-promoter interactions. Extensive studies have demonstrated that chromatin loops play an essential role in maintaining the 3D structure of the genome and facilitating gene regulation^22–25^.

The functional importance of chromatin loops makes it important to develop loop callers tailored to different input experimental datasets. All existing loop callers are designed for genomic data generated from proximity-ligation assays, which utilize the *count*-based statistical framework to model chromatin contact frequency^26^. As a result, these tools are inherently not directly applicable to imaging data, which allow the *continuous* measurement of Euclidean distances between targeted genomic segments of interest. To the best of our knowledge, no method is available to identify chromatin loops from such imaging data. In the wake of rapid development of chromatin tracing technologies, tailored computational methods to reliably identify chromatin loops from imaging data have become more critical. Importantly, such loop analysis methods may advance our understanding of the relationships between genome structure and gene regulation.

## 2. Results

### 2.1 SnapFISH algorithm

To fill in the abovementioned methodological gap, here we present Single-Nucleus Analysis Pipeline for multiplexed DNA FISH data (SnapFISH). SnapFISH takes multiplexed DNA FISH single bright spot localization data as input, and outputs the predicted chromosomal location of chromatin loops. Briefly, SnapFISH first collects the 3D localization coordinates of each genomic segment targeted by FISH (hereafter referred to as targeted segment) in each cell (**Figure 1A**), and computes the pairwise Euclidean distances between all imaged targeted loci (**Figure 1B**). Next, for all pairs of targeted segments found within a 1D genomic distance range of 100Kb ~ 1Mb, SnapFISH compares the pairwise Euclidean distances between the pair of interest and its local neighborhood region (**Figure S1**, see details in **Methods**) using a two-sample T-test (**Figure 1C**). SnapFISH then converts the resulting P-values into false discovery rates (FDRs), and defines a pair of targeted segments as a loop candidate if its T-test statistic is less than −4, FDR is less than 10% and the ratio between the average Euclidean distance in the local genomic neighborhood region and the average Euclidean distance at the pair of targeted segments of interest is above a pre-specified threshold (**Figure 1D**, see details in **Methods**). Lastly, SnapFISH groups nearby loop candidates into clusters, identifies the pair with the lowest FDR within each cluster (hereafter referred as the cluster summit), and uses these summits as the final list of chromatin loops (**Figure 1D**, see details in **Methods**).

**Figure 1.**
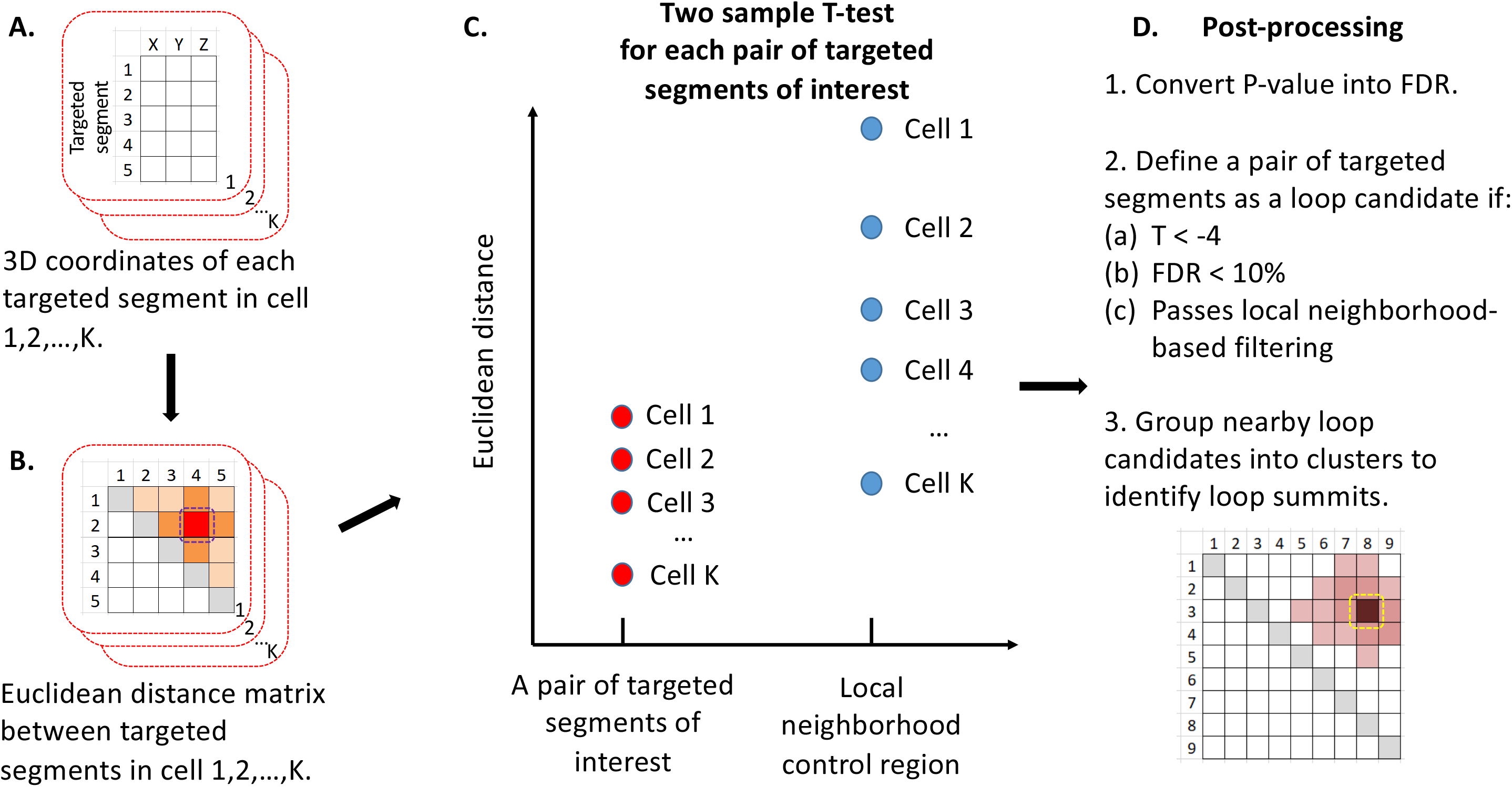
The flowchart of SnapFISH algorithm. **A.** The 3D coordinates for each targeted segment in cell 1, 2,…, *K*. Each matrix represents one cell. Each row is one targeted segment and *X, Y, Z* are the 3D coordinates. **B.** Euclidean distance matrices. Again, each matrix is for one cell. The dashed purple block highlighted the pair of targeted segments of interest: between the targeted segment 2 and the targeted segment 4. **C.** Two-sample T-test comparing the Euclidean distance between the pair of targeted segments of interest and its local neighborhood control region. **D.** Post-processing to identify loop candidates and loop summits. The cartoon represents the clustering of nearby loop candidates and the selection of loop summit (i.e., between the targeted segment 3 and the targeted segment 8), which is highlighted by the dashed yellow block. Detailed definition of local neighborhood regions is listed in **Methods**.

### 2.2 Applying SnapFISH to multiplexed DNA FISH data and ORCA data in mESCs at the *Sox2* locus

To demonstrate the accuracy and reliability of SnapFISH, we first applied SnapFISH to a multiplexed DNA FISH dataset previously generated from mouse embryonic stem cells (mESCs) to investigate the chromatin conformation at the *Sox2* locus^16^ (such data is downloaded from the 4D Nucleome data portal, see details in **imaging data resource**). The 205Kb chromosomal target region (mm10: chr3:34,601,078-34,806,078) imaged in the experiments spans both the promoter of the *Sox2* gene and its super-enhancer, which is located ~100Kb downstream. Previous studies have identified a chromatin loop between the *Sox2* promoter and its super-enhancer via 5C^27^, and validated the functional importance of such loop via the CRISPR experiment, which showed that deletion of the super-enhancer reduce >90% of the *Sox2* expression^28^. The mESCs used in the original study^16^ have two haplotypes: the CAST allele and the 129 allele. Notably, the CAST allele contains a 7.5Kb insertion containing 4 CTCF-binding sites (hereafter referred to as 4CBS) between the *Sox2* promoter and its super-enhancer, while the 129 allele does not contain the insertion. The multiplexed DNA FISH experiments involved 41 probe sets, each corresponding to individual 5Kb genomic target segments tiling the 205Kb region of interest. In addition, there was one extra probe for the 7.5Kb insertion on the CAST allele, which permits the differential identification of the CAST allele and the 129 allele in the same nucleus. Previous study^16^ showed that the CAST allele-specific 7.5Kb 4CBS insertion resulted in weaker chromatin looping strength between the *Sox2* promoter and its super-enhancer.

We re-analyzed the multiplexed DNA FISH data (see details in **Methods**) from a total of 1,416 cells (i.e., 1,416 CAST alleles and 1,416 129 alleles) using SnapFISH. The average targeted segment detection efficiency, defined as the proportion of imaged targeted segments among all targeted segments, is 71.9% and 69.3% for the CAST allele and the 129 allele, respectively. We first computed the average Euclidean distances between all pairs of 5Kb genomic targeted segment in these experiments (**Figure S2A**), and compared that with the 5Kb bin resolution chromatin contact frequency in the mESC bulk Hi-C data^16^ in an allele-specific manner (**Figure S2B**). As we expected, the average spatial distance measured from multiplexed DNA FISH data is closely correlated with the Hi-C contact frequency. The Pearson correlation coefficient between Hi-C contact frequency and the inverse of average Euclidean distances is 0.812 and 0.805 for the CAST allele and the 129 allele, respectively. Applying SnapFISH to 1,416 CAST alleles and 1,416 129 alleles identified 4 and 13 loop candidates, respectively (**Figure S2C**). Finally, SnapFISH grouped neighboring loop candidates, and detected a single chromatin loop summit between the *Sox2* promoter and the super-enhancer in both CAST and 129 alleles (**Figure 2A**). As a comparison, we applied HiCCUPS to identify 5Kb bin resolution chromatin loops from mESC bulk Hi-C data^29^, and found that SnapFISH-identified loop are also near the HiCCUPS loop (**Figure 2A**).

**Figure 2.**
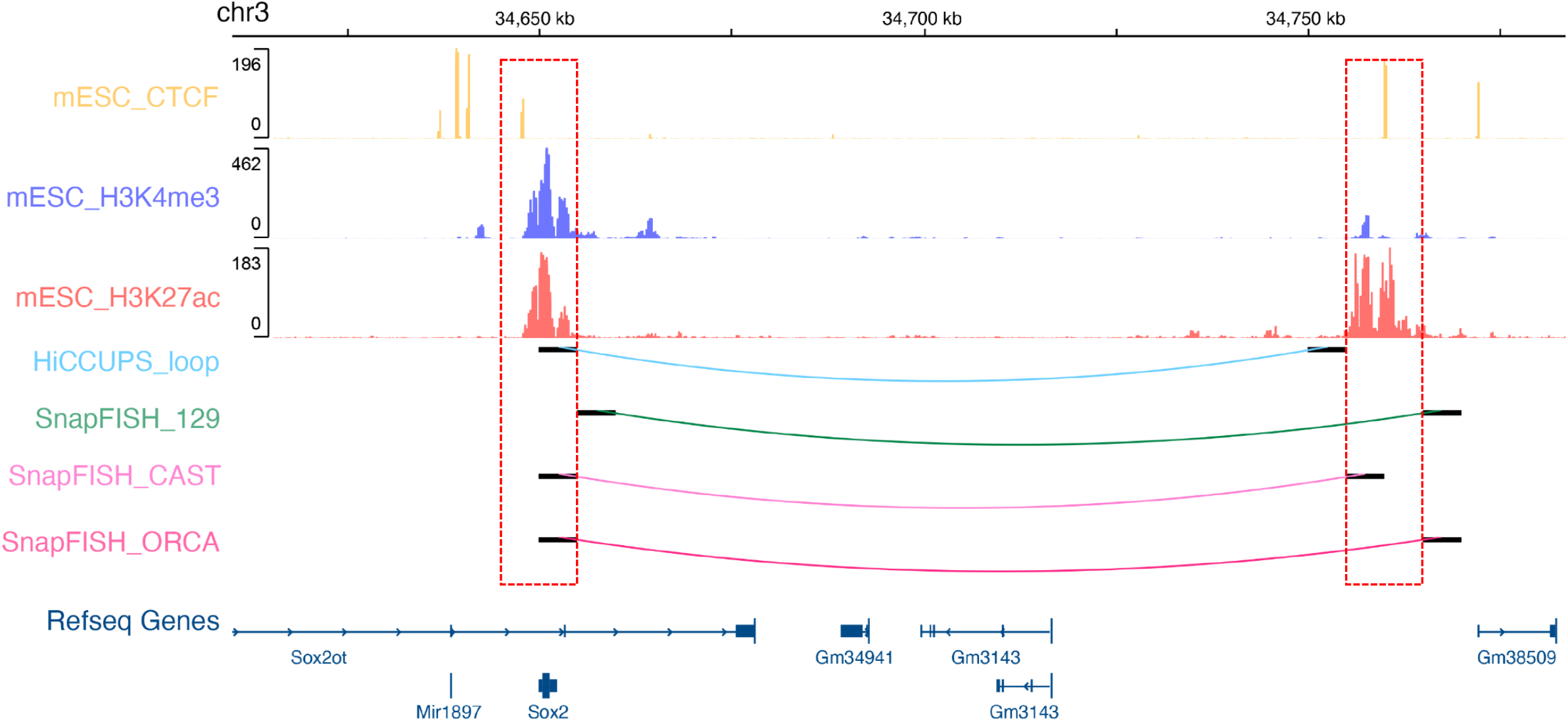
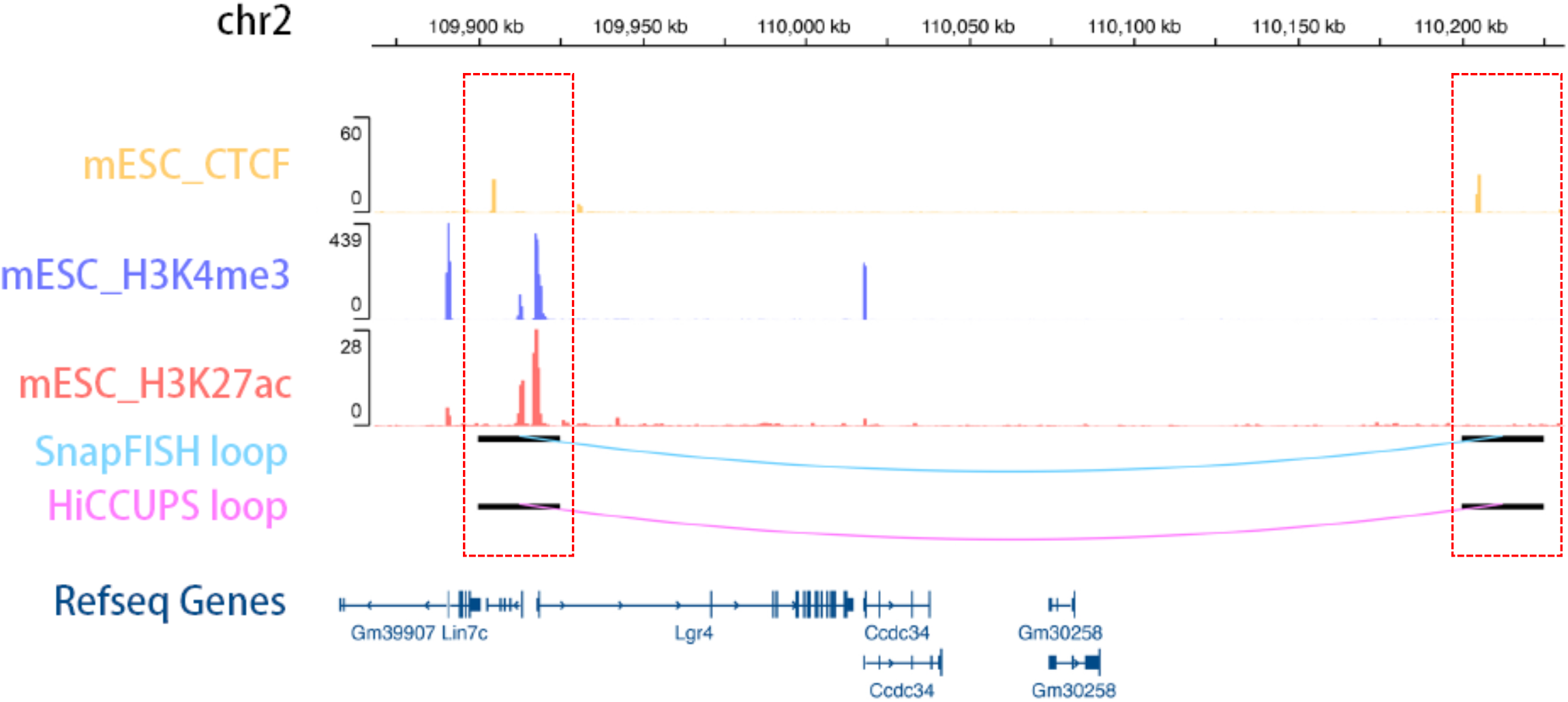
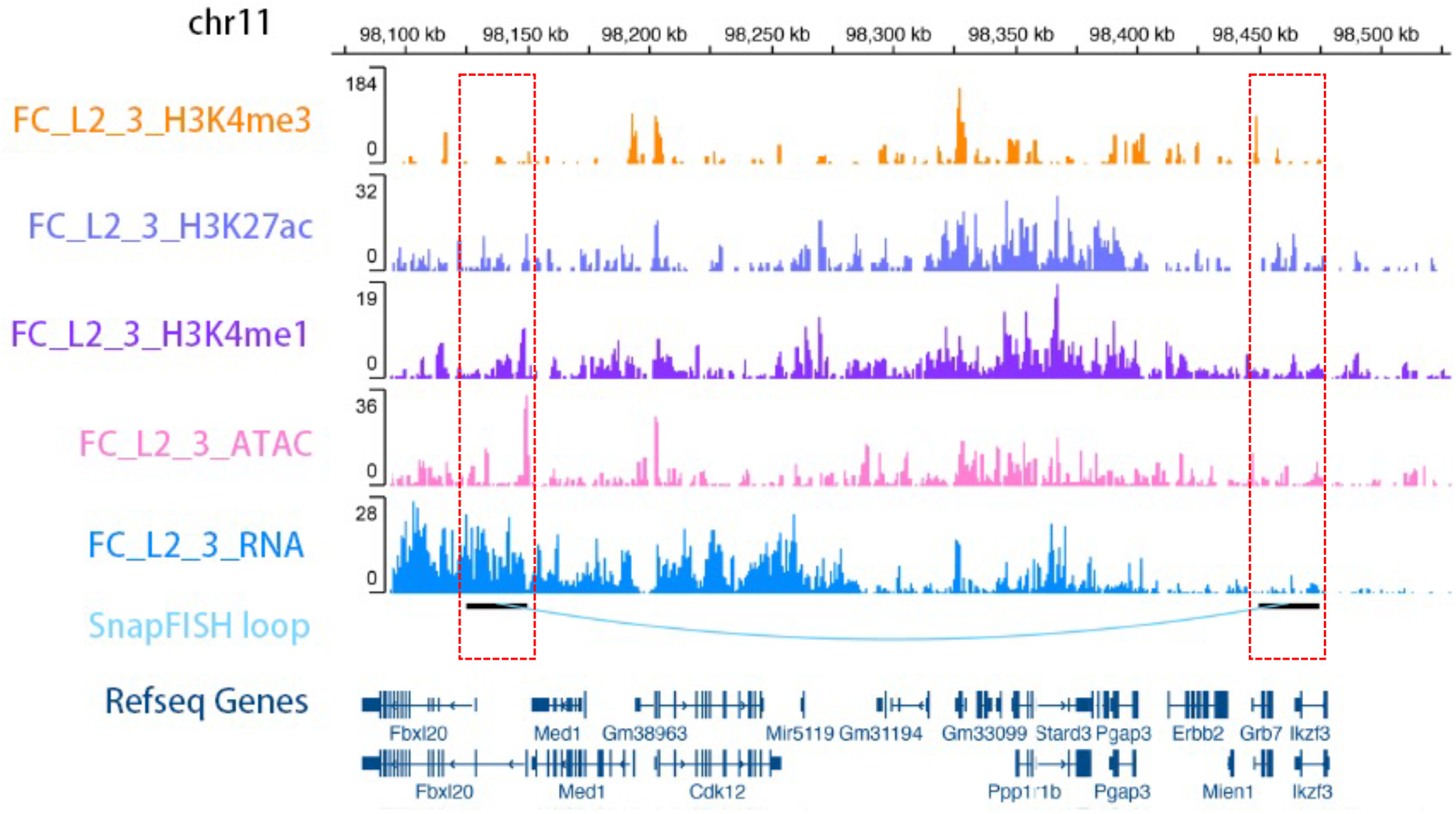
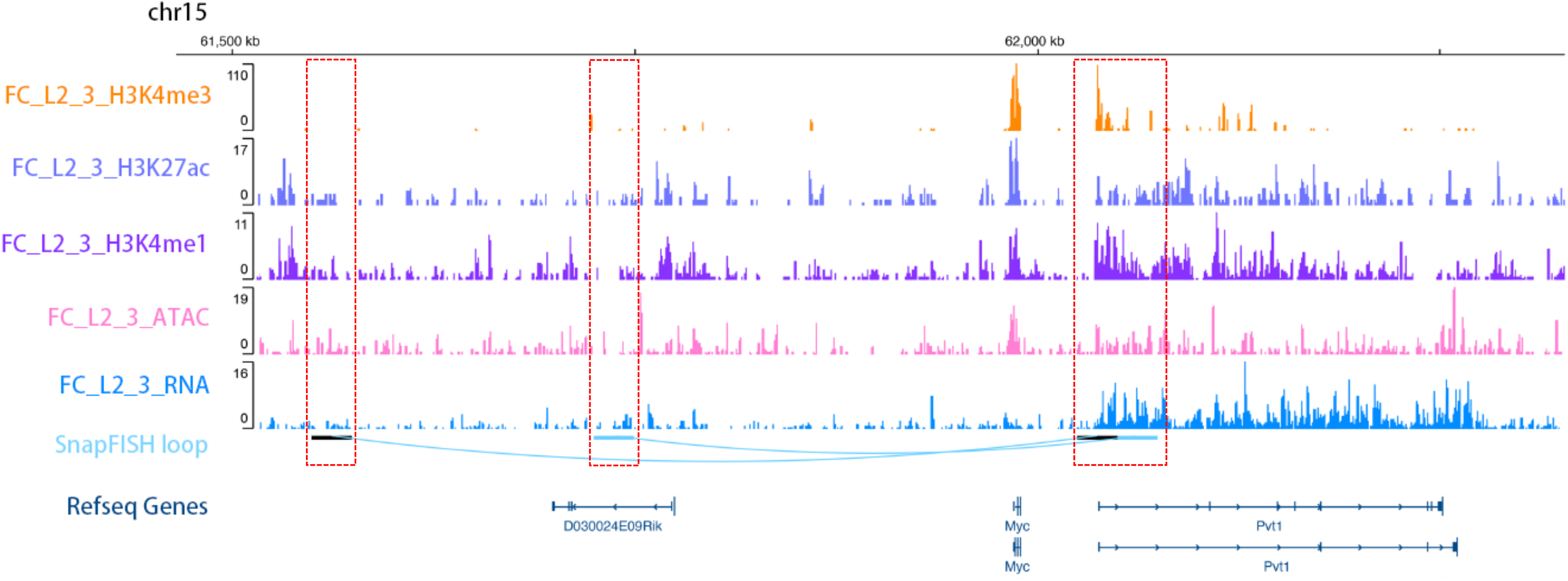
SnapFISH identified chromatin loops with high accuracy. **A.** SnapFISH identified the *Sox2* enhancer-promoter loop from mESC multiplexed DNA FISH data and ORCA data. The top three tracks are mESC CTCF, mESC H3K4me3^33^, and mESC H3K27ac ChIP-seq data. The middle four arcs represent the HiCCUPS-identified loop from mESC bulk Hi-C data^29^, the SnapFISH-identified loop from 1,416 129 alleles and 1,416 CAST alleles, and the SnapFISH-identified loop from mESC ORCA data, respectively. All these loops are at 5Kb bin resolution. The bottom track is the Refseq gene annotation. The red dashed box on the left and the right represents the location of the promoter of *Sox2* gene and its super-enhancer, respectively. We allow for 5Kb gap between loop anchors and the *Sox2* promoter or super-enhancer (see details in **Methods**). **B.** SnapFISH identified a CTCF-CTCF loop in chr2 from mESC DNA seqFISH+ data. The top three tracks are mESC CTCF, mESC H3K4me3 and mESC H3K27ac ChIP-seq data. The middle two tracks are the SnapFISH-identified loop and the HiCCUPS-identified loop, both at 25Kb bin resolution. The bottom track is the Refseq gene annotation. Both anchors (dashed red boxes) contain mESC CTCF ChIP-seq peaks. **C.** SnapFISH identified a promoter-promoter loop in chr11 from DNA seqFISH+ data in mouse excitatory neurons. The top five tracks are H3K4me3, H3K27ac, H3K4me1, ATAC-seq and RNA-seq data from cortical excitatory neurons L2/3 collected from mouse frontal cortex tissue (FC_L2_3 for short)^30^. The bottom two tracks are the SnapFISH-identified 25Kb bin resolution loop and the Refseq gene annotation. The left anchor (dashed red box on the left) is at the promoter of gene *Fbxl20*. The right anchor (dashed red box on the right) is at the promoter of gene *Grb7*. **D.** SnapFISH identified two enhancer-promoter loops in chr15 from DNA seqFISH+ data in mouse excitatory neurons. The top five tracks are H3K4me3, H3K27ac, H3K4me1, ATAC-seq and RNA-seq data from cortical excitatory neurons L2/3 collected from mouse frontal cortex tissue (FC_L2_3 for short)^30^. The bottom two tracks are the SnapFISH-identified 25Kb bin resolution loop and the Refseq gene annotation. The right anchor (dashed red box on the right) is at the promoter of gene *Pvt1*. The two left anchors (two dashed red boxes on the left and in the middle) both contain active enhancer mark H3K4me1 and H3K27ac peaks and open chromatin mark ATAC peaks.

Next, we evaluated the sensitivity of SnapFISH using different numbers of input target alleles. Specifically, we ranked all 1,416 CAST alleles by their targeted segment detection efficiency, and selected the top 1,400, 1,200, 1,000, 800, 600, 400, 200, 100 and 50 CAST alleles as input for SnapFISH. We performed the same analysis with the 129 allele. Consistent with previous findings^16^ indicating that the strength of *Sox2* enhancer-promoter loop is weaker in the CAST allele compared to that in the 129 allele, SnapFISH found fewer loop candidates among the CAST allele than that in the 129 allele (**Table S1**), due to the CAST-specific 4CBS insertion. However, despite these differences, we observed that SnapFISH can accurately identify the *Sox2* enhancer-promoter loop with as few as 200 cells, for both CAST allele and 129 allele (**Figure S3A** and **Figure S3B**). Our results suggest that SnapFISH achieves high sensitivity even with a small number of cells.

Since not all targeted segments can be detected in the multiplexed DNA FISH data, we further evaluated how targeted segment detection efficiency affects the sensitivity of SnapFISH. Specifically, we again ranked all 1,416 CAST alleles by their targeted segment detection efficiency, and then equally divided them into three groups, where each group consisted of 472 CAST alleles. We stratified them as “high”, “median” and “low” targeted segment detection efficiency group, with average targeted segment detection efficiency 85.4%, 73.2% and 57.0%, respectively. We performed the same analysis for the 1,416 129 alleles, and similarly created three 129 allele groups, with average targeted segment detection efficiency 82.6%, 70.4% and 55.0%, respectively. We applied SnapFISH to each of the six groups and examined the identified loop candidates and loop summits. In the CAST alleles where the *Sox2* enhancer-promoter loop strength is weak due to the CAST-specific 4CBS insertion, SnapFISH can only detect the loop in the “high” targeted segment detection efficiency group (**Figure S3C** and **Table S2A**). In contrast, in the 129 alleles, SnapFISH can detect the loop in both “high” and “median” targeted segment detection efficiency groups (**Figure S3C** and **Table S2B**). Notably, in the 129 alleles, while the SnapFISH-identified loop from the “high” targeted segment detection efficiency group overlaps the superenhancer, the one identified from the “median” targeted segment detection efficiency group is 5Kb away from the super-enhancer (**Figure S3C**). Such deviation is probably due to higher level of uncertainty caused by the larger proportion of missing targeted segments. As expected, our results suggest that targeted segment detection efficiency can influence the sensitivity of detecting chromatin loops.

Additionally, we evaluated the performance of SnapFISH on an ORCA dataset^10^ generated at the same *Sox2* locus in mESCs (see details in **imaging data resource**). Specifically, Mateo et al imaged the 170Kb region around the *Sox2* gene (mm10: chr3:34,601,078-34,771,078) in mESCs, which consists of 34 5Kb targeted segments. Across all 6,007 imaged alleles, the average targeted segment detection efficiency is 56.9%. Consistent with the results in multiplexed DNA FISH data, SnapFISH identified the *Sox2* enhancer-promoter loop (**Figure 2A**). Taken together, our data show that SnapFISH can accurately identify the *Sox2* enhancer-promoter loop, which had previously been identified by mESC bulk Hi-C data, from both multiplexed DNA FISH data and ORCA data.

### 2.3 Applying SnapFISH to DNA seqFISH+ data in mESCs and mouse excitatory neurons

Encouraged by the results from multiplexed DNA FISH data and ORCA data in mESCs at the *Sox2* locus, we then applied SnapFISH to a recently published DNA seqFISH+ dataset^8^. We first re-analyzed a publicly available DNA seqFISH+ dataset from mESCs^8^, where the authors selected one region from each chromosome, with region size ranging 1.5Mb ~ 2.35Mb (**Table S3**), and performed DNA seqFISH+ experiment at 25Kb bin resolution, in two biological replicates. We combined data from both biological replicates, resulting in 446 cells (i.e., 892 alleles) in total. The average targeted segment detection efficiency is 65.2%. Note that in DNA seqFISH+ data, one cannot distinguish the paternal allele and the maternal allele in the same cell. We applied SnapFISH to detect chromatin loops from autosomes, with 1D genomic distance ranging from 100Kb ~ 1Mb, and identified 6 chromatin loops (**Table S4A**). To validate the results, we analyzed a deeply sequenced mESC bulk Hi-C data^29^, and identified 16 loops with HiCCUPS at 25Kb resolution in the corresponding genomic region where DNA seqFISH+ data is available (**Table S5**). We found that all six SnapFISH identified chromatin loops overlap with HiCCUPS loops (**Figure 2B**, **Figure S4** and **Figure S5**). The precision and recall are 1.000 and 0.375, respectively. **Figure 2B** shows an illustrative example in chr2, where SnapFISH identified a CTCF-CTCF loop, which has also been detected by HiCCUPS from mESC bulk Hi-C data^29^. Taken together, our results show that when applying to mESC DNA seqFISH+ data, SnapFISH can accurately and reliably identify loops that were previously identified by mESC bulk Hi-C data.

We additionally re-analyzed another publicly available 25Kb resolution DNA seqFISH+ dataset from mouse cerebral cortex tissue^9^ (**Table S3**). The dataset also included RNA seqFISH data simultaneously generated from the same cells^9^, which can be used to group and annotate distinct cell clusters. We combined the three biological replicates of DNA seqFISH+ datasets to obtain a total of 2,762 cells (i.e., 5,524 alleles), with the average targeted segment detection efficiency of 40.9%. Due to the relatively low targeted segment detection efficiency, we only applied SnapFISH to the excitatory neurons^9^ consisting of 1,895 cells (i.e., 3,790 alleles, with targeted segment detection efficiency 43.1%), and identified 28 loops (**Table S4B** and **Figure S6**).

To the best of our knowledge, there is no publicly available bulk Hi-C data or single cell Hi-C data generated from excitatory neurons in mouse cerebral cortex tissue. To evaluate the functional relevance of these SnapFISH identified loops, we used a recent published Paired-Tag data^30^ to obtain transcriptomic data and epigenetic data from cortical excitatory neurons L2/3 collected from mouse frontal cortex tissue (FC_L2_3 for short), including active promoter mark H3K4me3, two active enhancer marks H3K27ac and H3K4me1, and the open chromatin region mark ATAC. **Figure 2C** shows an illustrative example of promoter-promoter loop in chr15, where the loop connects the promoter of gene *Fbxl20* and the promoter of gene *Grb7*. Both loop anchors overlap H3K4me3 peaks, H3K27ac and H3K4me1 peaks, and ATAC-seq peaks. In addition, **Figure 2D** shows two enhancer-promoter loops in chr16, where two anchors on the right are both near the transcription start site of gene *Pvt1*, and contain active promoter mark H3K4me3 peaks. The two anchors on the left both overlap H3K27ac, H3K4me1 peaks, and ATAC-seq peaks. We obtained similar results when applying SnapFISH to three additional mouse brain cell types (**Supplementary Information Section 1**). Taken together, our results suggest that SnapFISH is able to identify putative enhancer-promoter loops from DNA seqFISH+ data in mouse brain tissue sample.

## 3. Conclusion and discussion

In this work, we report SnapFISH, the first computational pipeline to identify chromatin loops from multiplexed DNA FISH data. We applied SnapFISH to multiplexed DNA FISH, ORCA and DNA seqFISH+ experiments in both mouse mESCs and mouse excitatory neurons, and benchmarked the performance of SnapFISH-identified chromatin loops using chromatin loops identified from bulk Hi-C data. We also showed the high reproducibility of SnapFISH between biological replicates (**Supplementary Information Section 2**). Additionally, SnapFISH is computationally efficiency, with the computing time increases linearly with the number of cells (**Supplementary Information Section 3**).

Building upon these promising results, we envision at least three directions warranted further investigation. First of all, the sensitivity of SnapFISH is can be further improved, in particular when applying to DNA seqFISH+ data in mESCs. As expected and as we show in the analysis of multiplexed DNA FISH data in mESCs (**Figure S3C** and **Table S2**), low level of targeted segment detection efficiency can reduce the sensitivity of loop detection. We expect that imputing missing 3D coordinates and missing Euclidean distance between genomic loci of interest may increase the sample size for the two sample T-test used in the SnapFISH algorithm (**Figure 1C**), and help to enhance statistical power of loop detection.

In addition, we only considered pair-wise chromatin interactions in this work. Multiplexed DNA FISH data provide rich information on multi-way chromatin interactions, making it feasible to detect events where one enhancer interacts with multiple target genes, or one gene’s promoter interacts with multiple enhancers simultaneously. We will extend our SnapFISH framework to identify multi-way chromatin interactions, and benchmark our findings with data generated from orthogonal technologies, including immunoGAM^31^ and scSPRITE^32^.

Last but not least, other genomic data modalities, including transcriptome and epigenome, can be imaged together with DNA in the same cell^8–10^. Integrating chromatin loops identified from SnapFISH with other genomic data modalities at single cell resolution, and characterizing their cell-to-cell variability, have the potential to reveal novel mechanisms of transcriptional regulation.

In summary, we developed SnapFISH, the first computational pipeline to identify chromatin loops from multiplexed DNA FISH data. As high-resolution multiplexed DNA FISH data are increasingly available, we consider SnapFISH a valuable tool in analyzing such data, facilitating better understanding of genome structure and genome function.

## Acknowledgements

We thank 4D Nucleome consortium investigators for comments and suggestions on the early version of this work. This study was funded by the NIH grants R35HG011922 (to M.H.) and U01DA052713 (to Y.L.). Y.L. was also partially funded by the NIH grants R01MH125236, R01HL163972, U01HG011720, and U24AR076730.

## Declaration of competing interest

B.R. is a cofounder and shareholder of Arima Genomics, Inc. and Epigenome Technologies, Inc. The remaining authors declare no competing interest.

## Reference genomes

We used mm10 for imaging data generated from mESCs and mouse excitatory neurons.

## Figure legends

**Figure S1.**
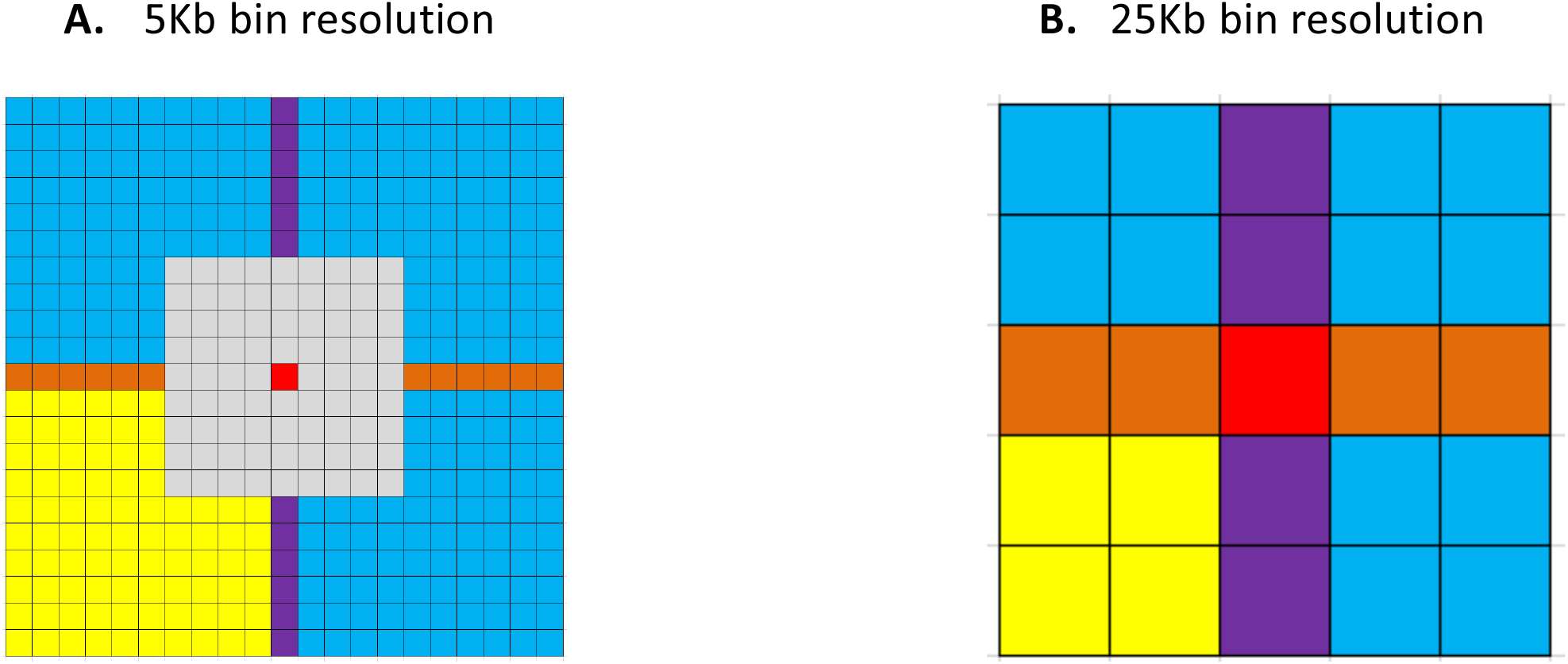
Cartoon illustration of local neighborhood control region. **A.** 5Kb bin resolution for multiplexed DNA FISH data and ORCA data. **B.** 25Kb bin resolution for DNA seqFISH+ data. In both, the red area represents the pair of targeted segments of interest. The blue, yellow, orange and purple areas represent the local neighborhood control region defined as a square with 25Kb~50Kb 1D genomic distance (hereafter referred to as square). Yellow areas represent the lower left region (hereafter referred to as LL). Purple areas represent the vertical region (hereafter referred to as V). Orange areas represent the horizontal region (hereafter referred to as H; see details in **Methods**). In both, blue areas represent the remaining part of the square. In **Figure S1A**, the grey areas are regions within 25Kb 1D genomic distance of the pair of targeted segments of interest, which are excluded from the definition of local neighborhood control region.

**Figure S2.**
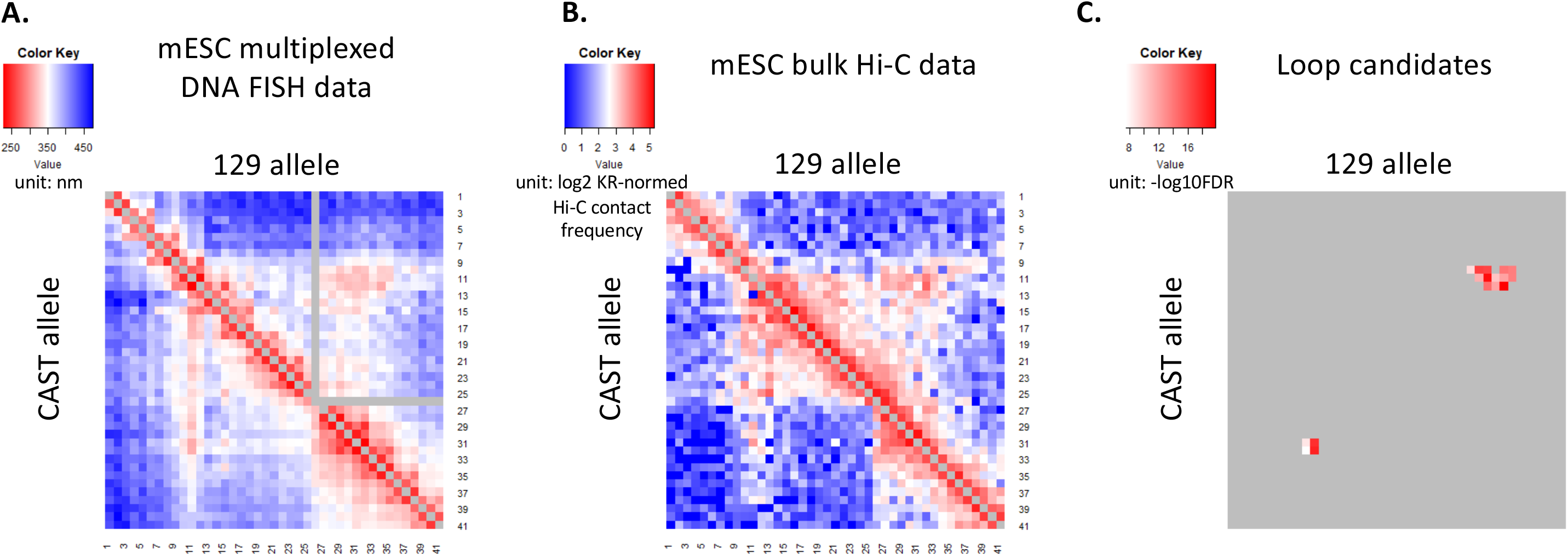
Applying SnapFISH to multiplexed DNA FISH data in mESCs. **A.** Average Euclidean distance (unit: nm) for all 5Kb resolution loci pairs calculated based on 1,416 CAST alleles (lower left panel) and 1,416 129 alleles (top right panel). Note that in 129 alleles, position #26 (chr3:34,725,000-34,730,000) is missing due to experimental error. **B.** 5Kb bin resolution log2 KR-normed Hi-C contact frequency matrix of mESC bulk Hi-C data^16^ (chr3:34.6Mb-34.805Mb) in the CAST allele (lower left panel) and the 129 allele (top right panel). **C.** Lower left panel: 4 loop candidates identified from 1,416 CAST alleles. Top right panel: 13 loop candidates identified from 1,416 129 alleles. The color is proportional to −log10 FDR at each loop candidate.

**Figure S3.**
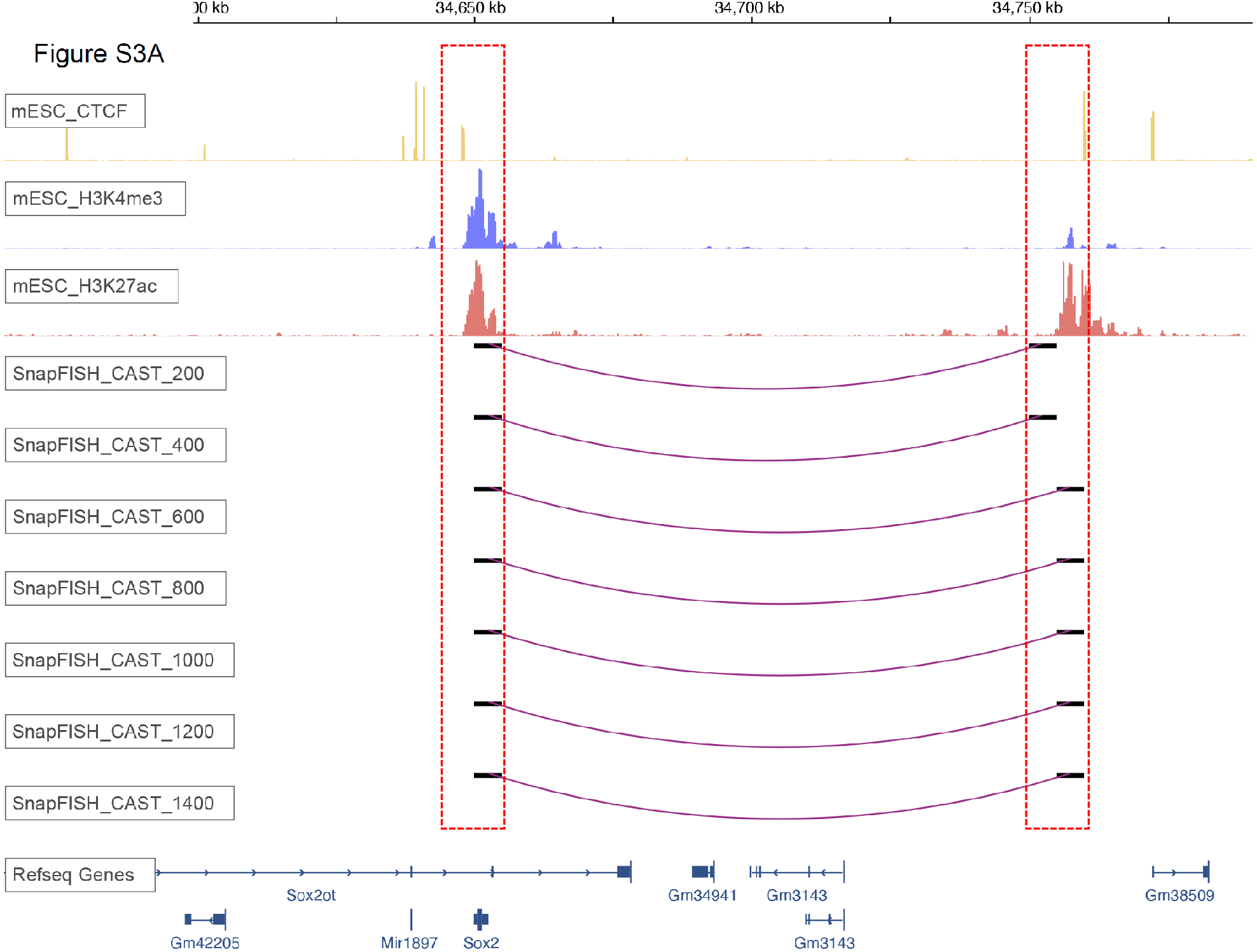
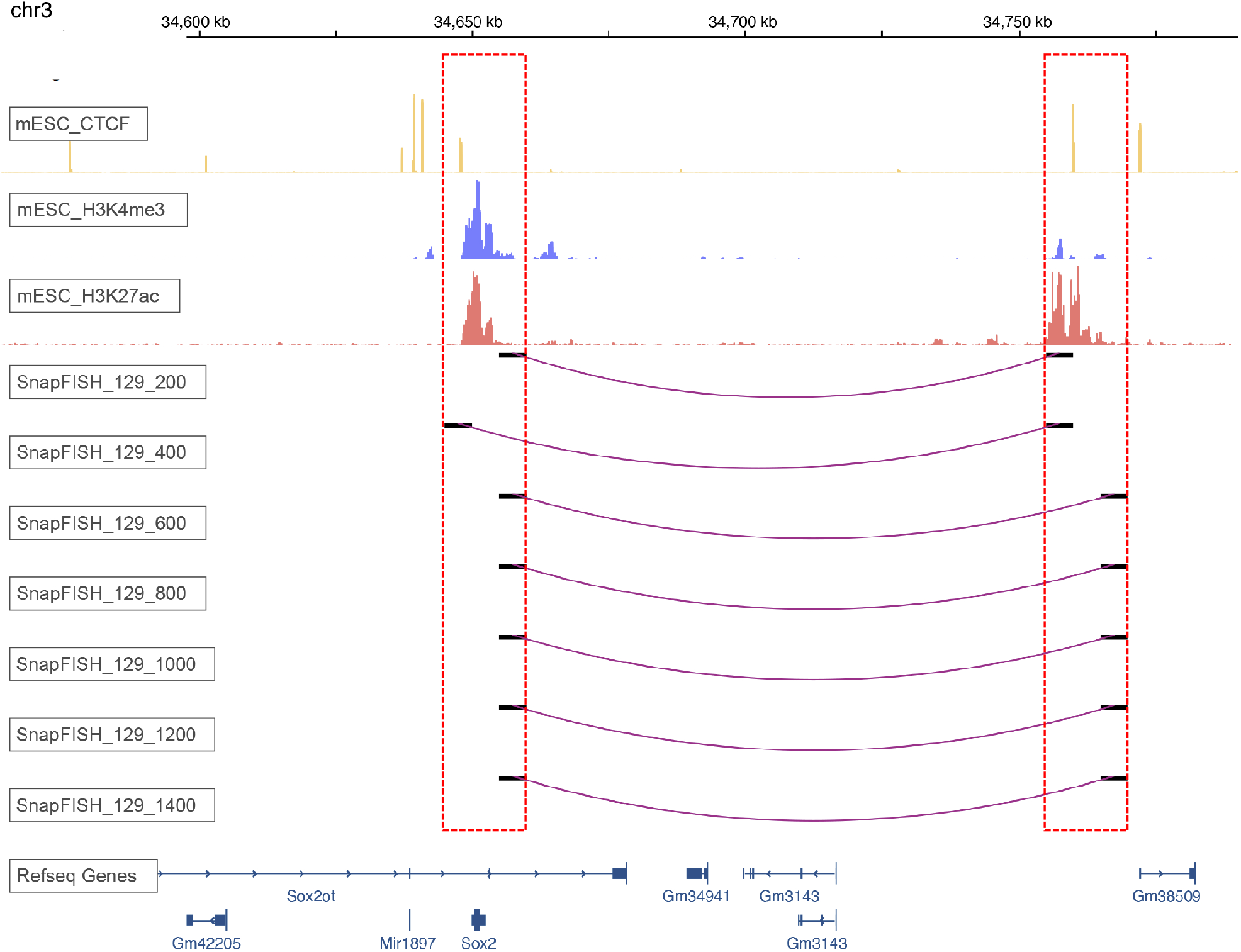
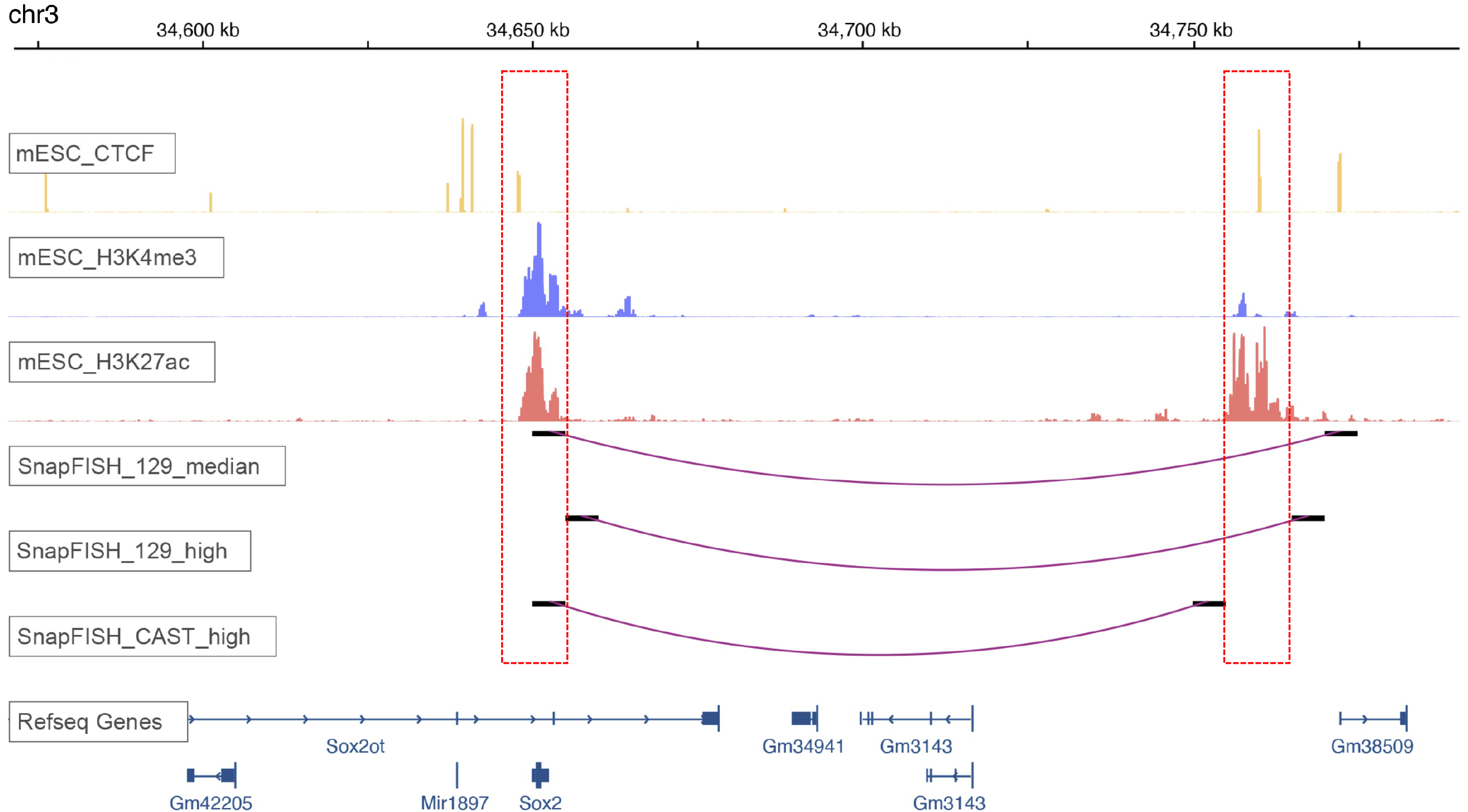
Benchmark the sensitivity of SnapFISH with different numbers of alleles and alleles with different levels of targeted segment detection efficiency. **A.** SnapFISH identified *Sox2* enhancer-promoter loop with more than 200 CAST alleles. **B.** SnapFISH identified *Sox2* enhancer-promoter loop with more than 200 129 alleles. **C.** SnapFISH identified *Sox2* enhancer-promoter loop with 472 CAST alleles or 472 129 alleles with three different levels of targeted segment detection efficiency.

**Figure S4.**
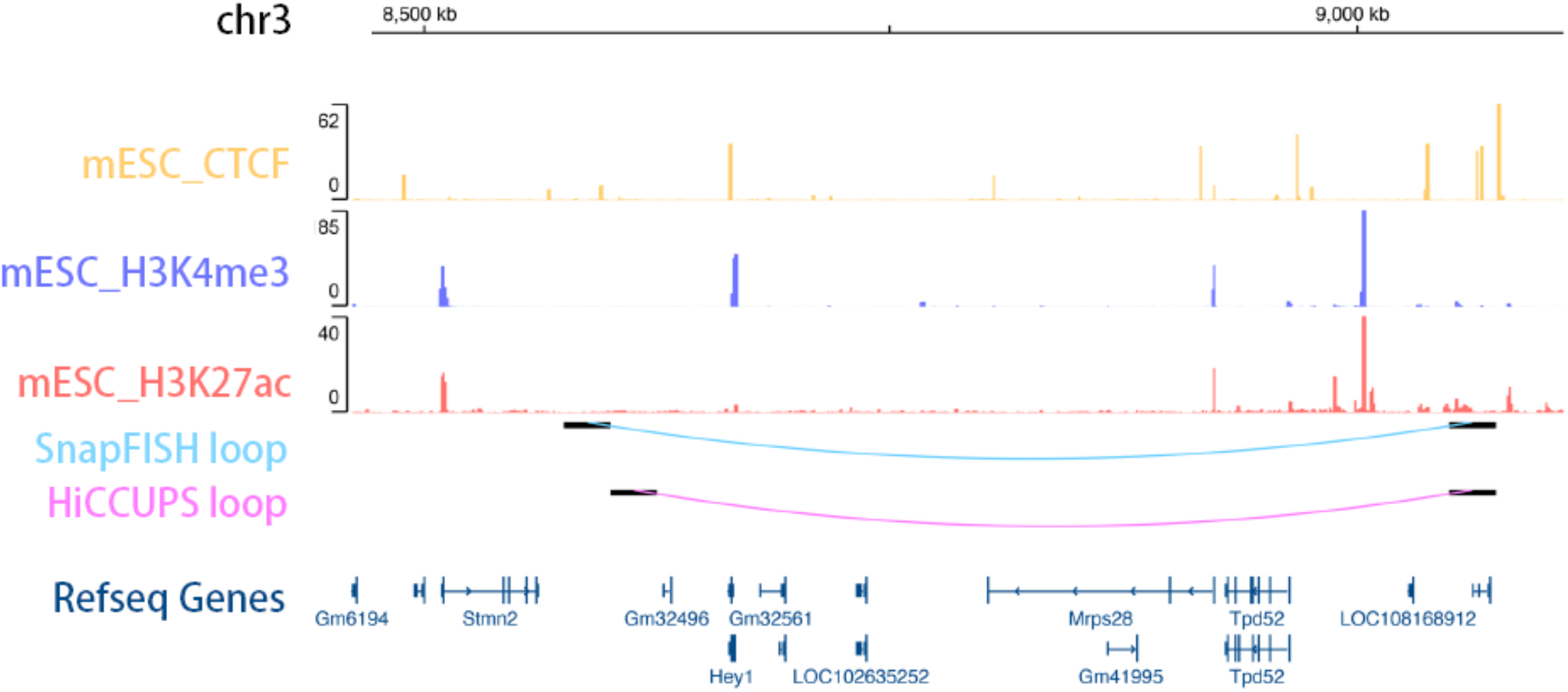
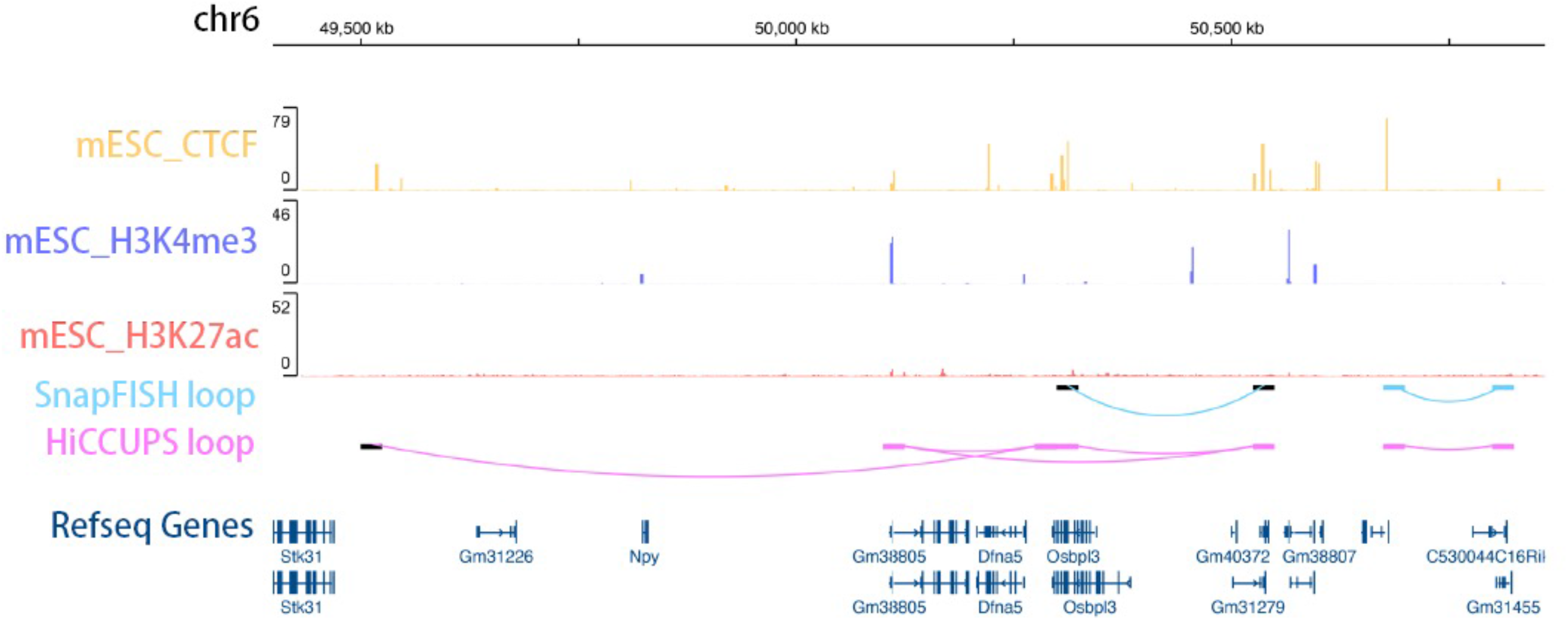
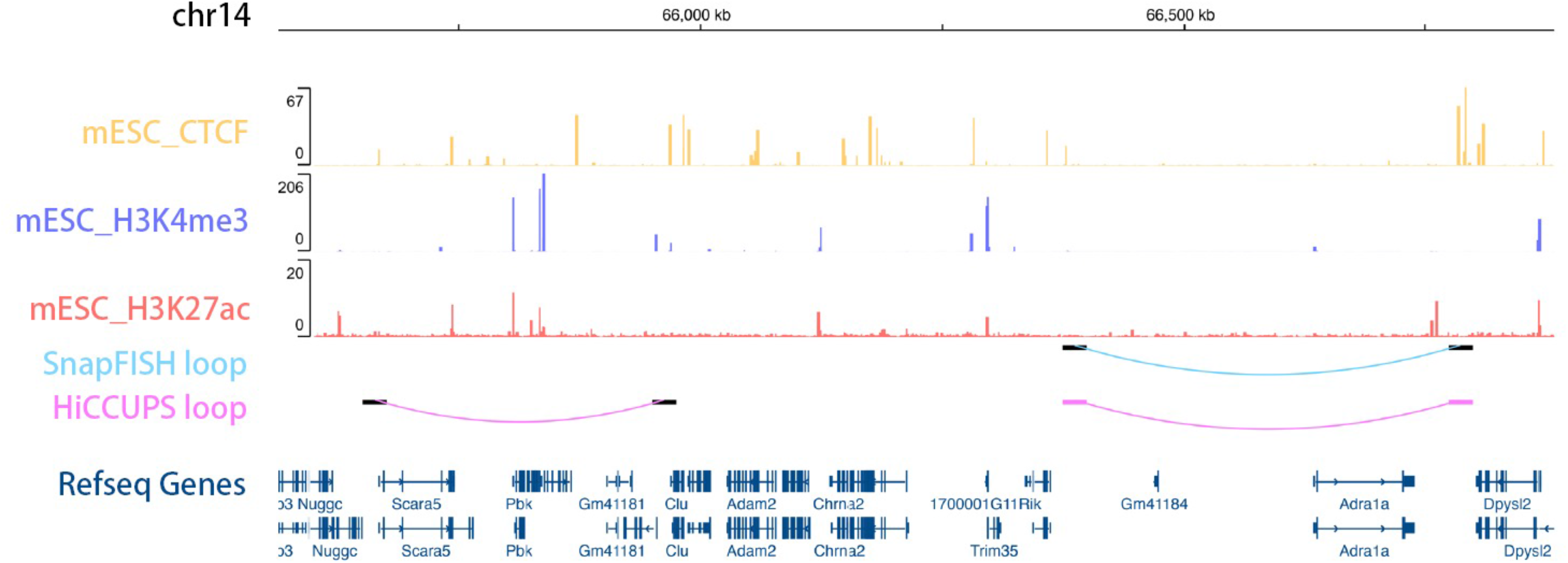
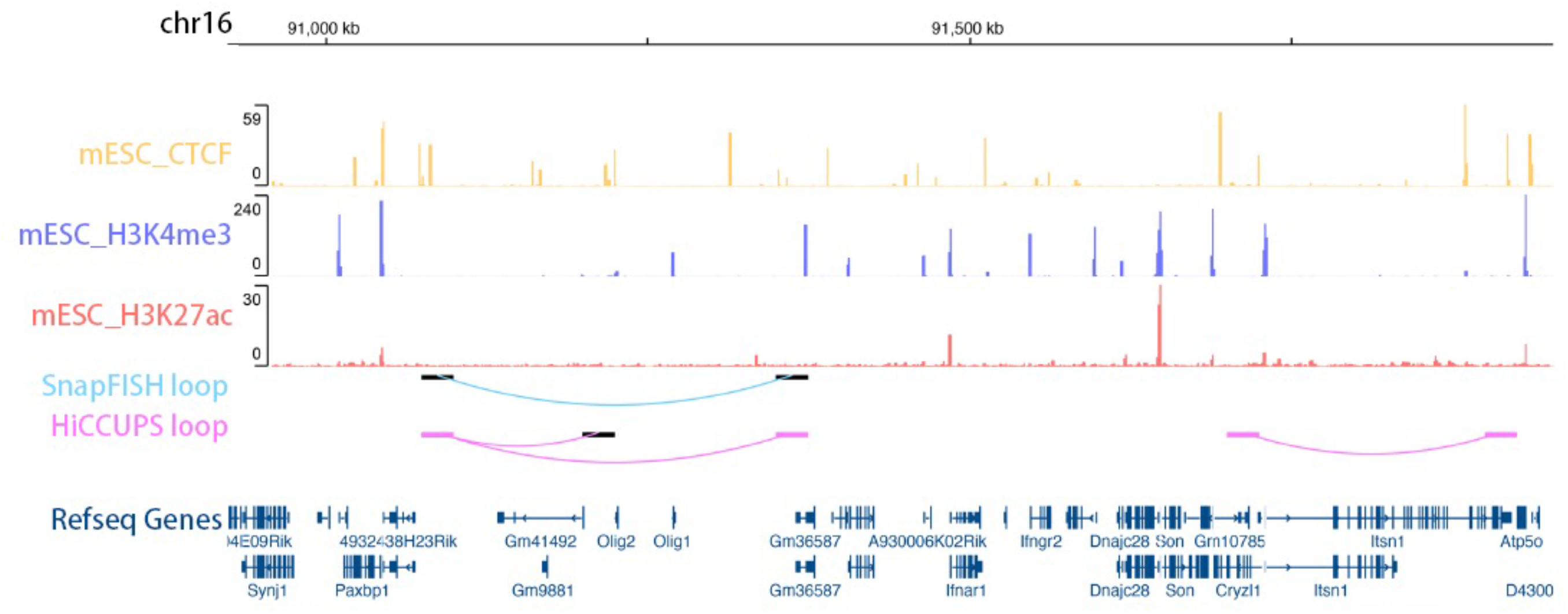
SnapFISH identified loops with high accuracy using DNA seqFISH data from mESC. Only chromosomes with SnapFISH-identified loops are showed. In each panel, the top three tracks are mESC CTCF, mESC H3K4me3 and mESC H3K27ac ChIP-seq data. The middle two tracks are the SnapFISH-identified loop and the HiCCUPS-identified loop, both at 25Kb bin resolution. The bottom track is the Refseq gene annotation. **A.** One SnapFISH-identified loop in chr3. **B.** Two SnapFISH-identified loops in chr6. **C.** One SnapFISH-identified loop in chr14. **D.** One SnapFISH-identified loop in chr16.

**Figure S5.**
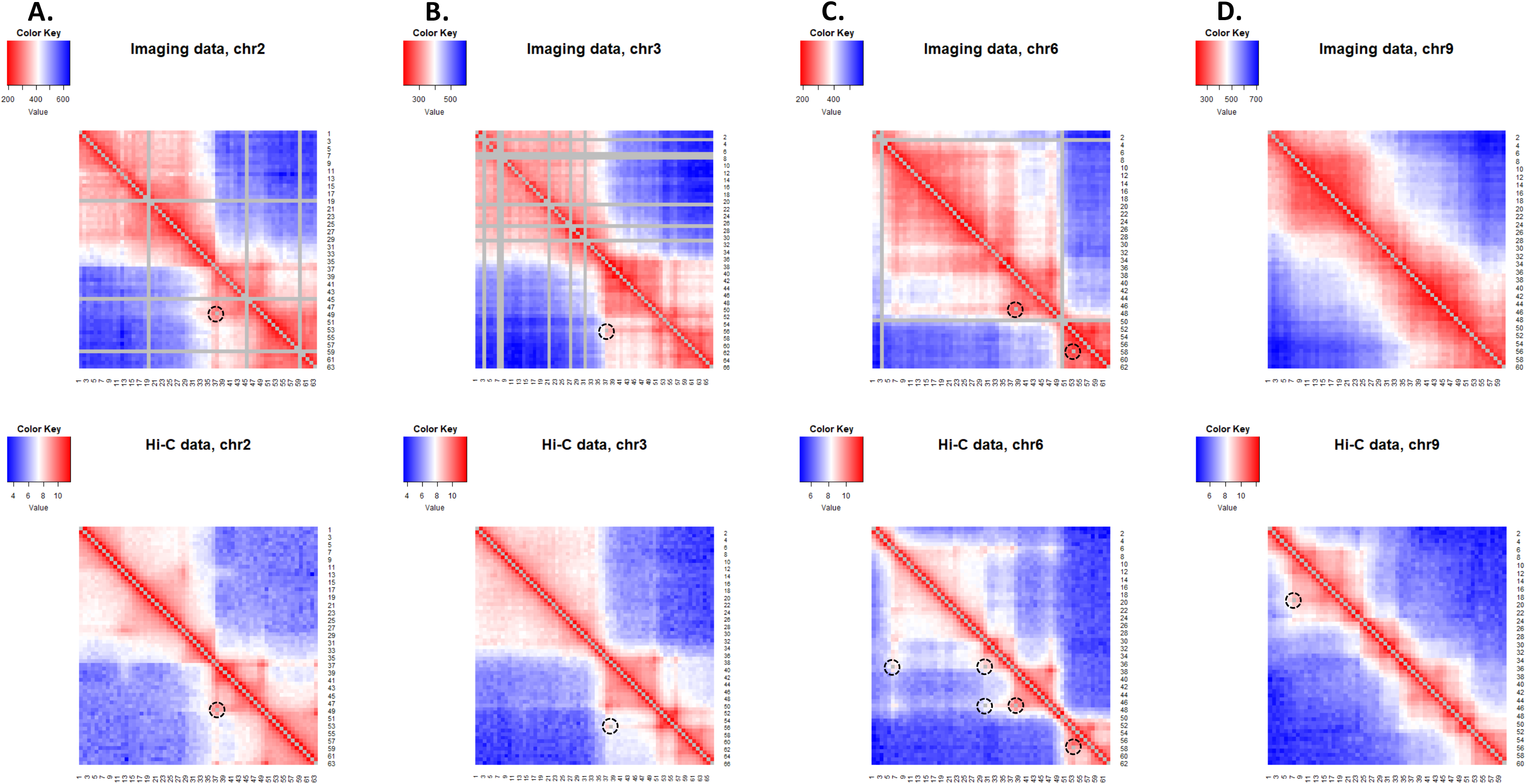
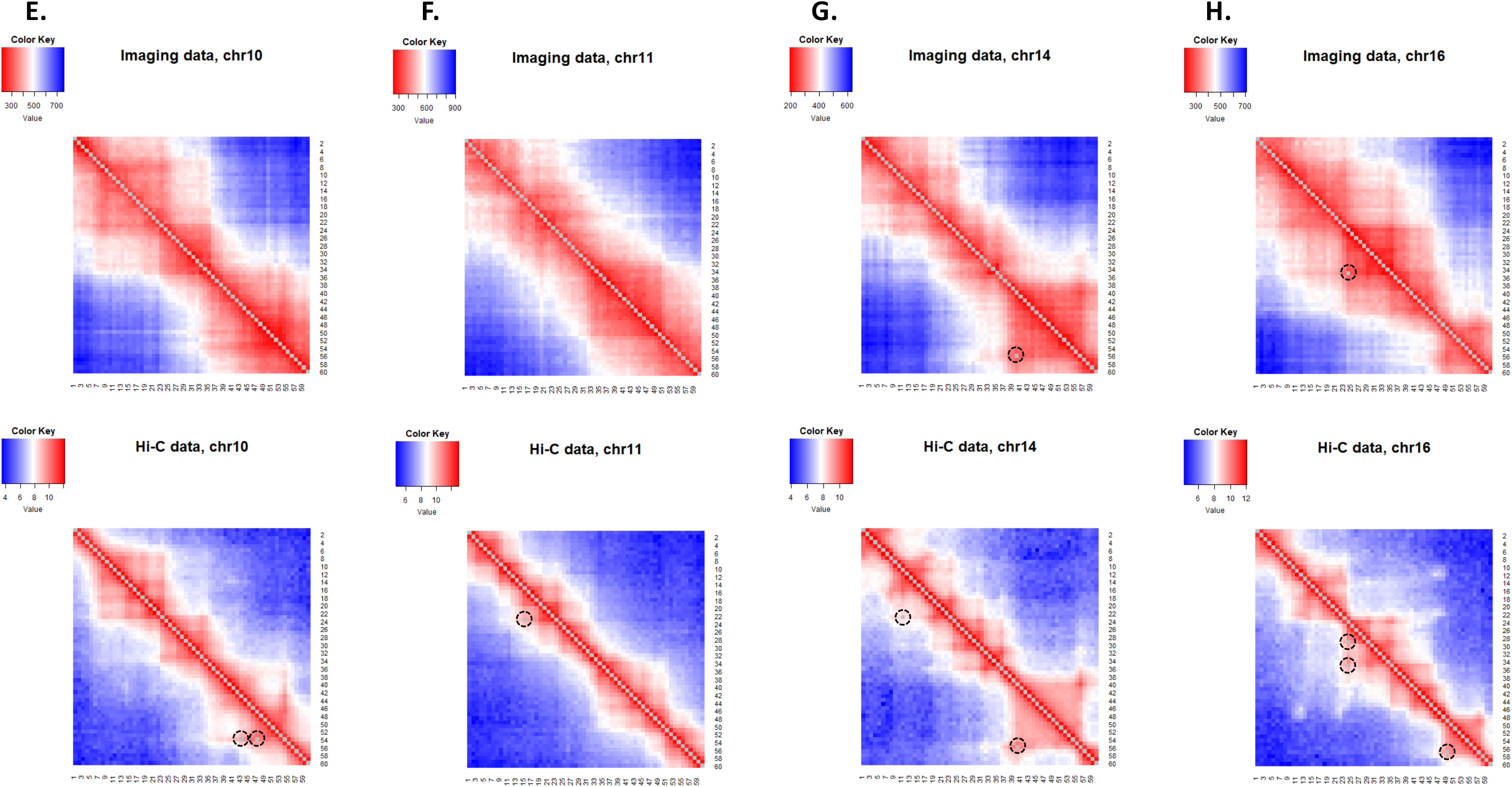
Heatmap of average Euclidean distance (unit: nm) from mESC DNA seqFISH+ data, and heatmap of Hi-C contact frequency (unit: log2 KR-normed Hi-C contact frequency) from mESC bulk Hi-C data. We showed eight chromosomes which contain 25Kb bin resolution HiCCUPS-identified loops: **A.** chr2. **B.** chr3. **C.** chr6. **D.** chr9. **E.** chr10. **F.** chr11. **G.** chr15. **H.** chr16. In each sub-figure, the top panel is the heatmap of average Euclidean distance from mESC DNA seqFISH+ data, and the bottom panel is the heatmap of Hi-C contact frequency from mESC bulk Hi-C data. Note that imaging data in chr2 (**A.**), chr3 (**B.**) and chr6 (**C.**) contain missing data, which are showed in grey vertical and horizontal lines. In each heatmap, the dashed black circles (with the grey pixel in the middle) in the lower left triangle represent SnapFISH-identified loops (top), or HiCCUPS-identified loops (bottom).

**Figure S6.**
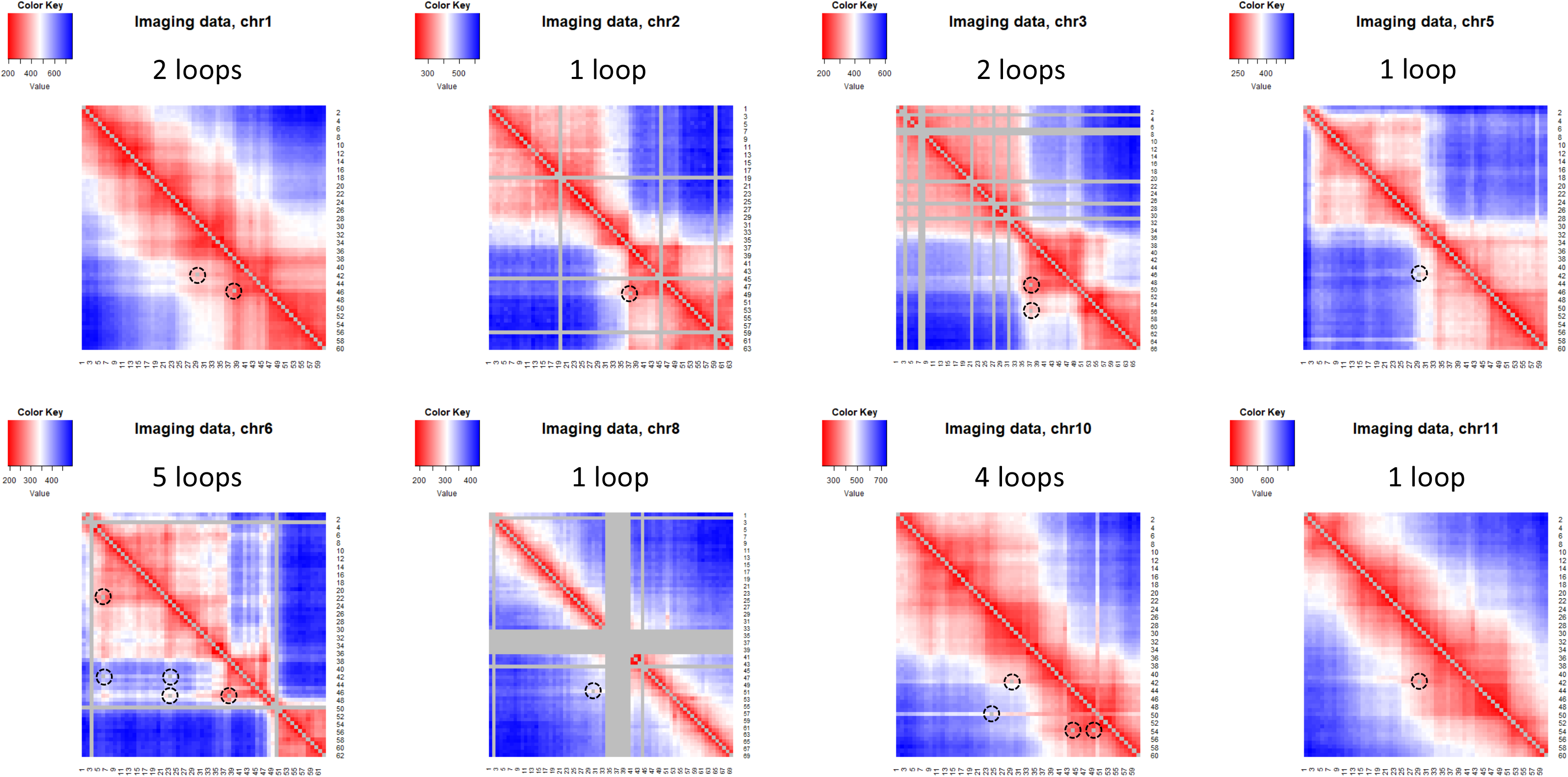
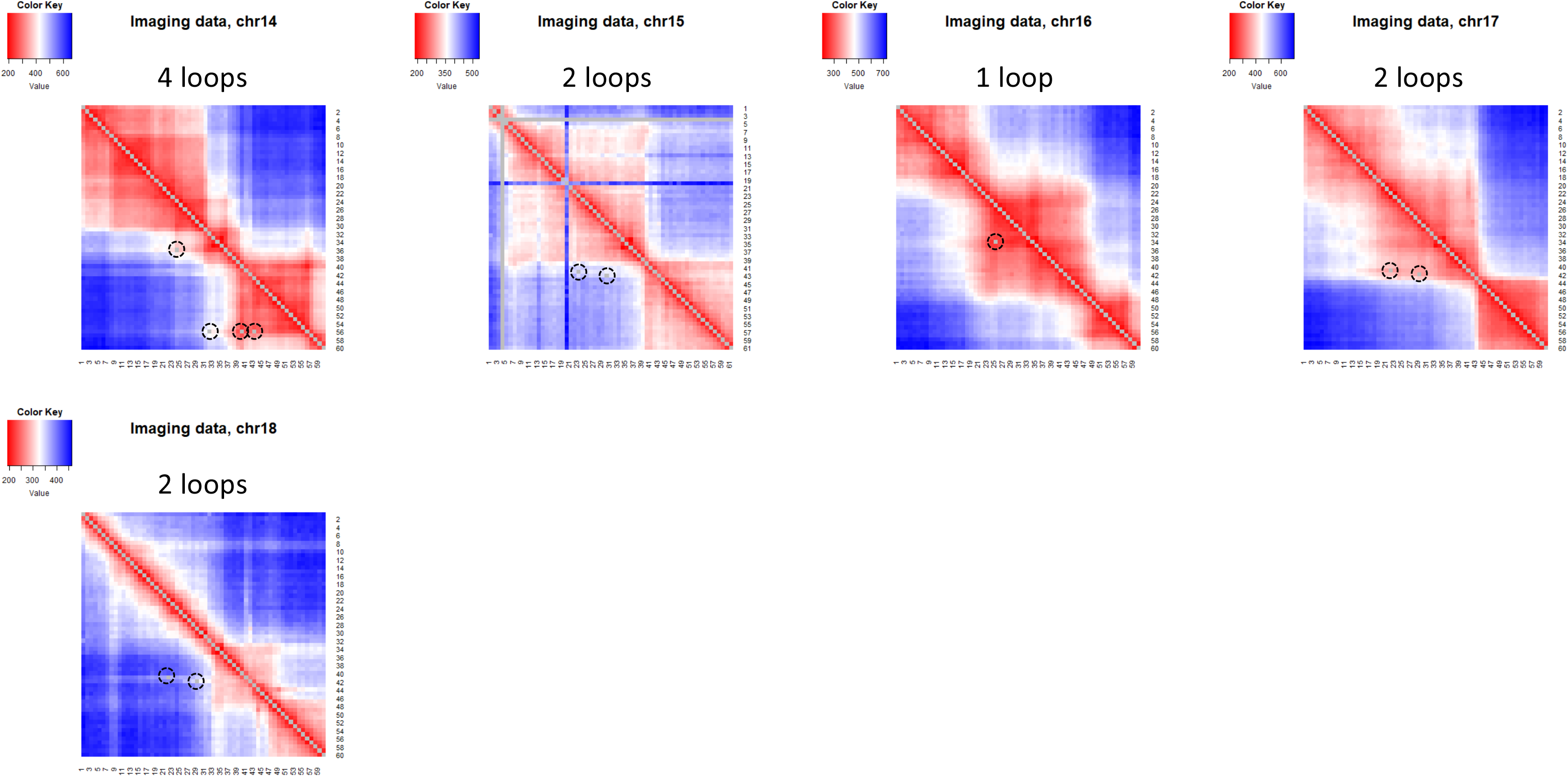
Heatmap of average Euclidean distance (unit: nm) from DNA seqFISH+ data in mouse excitatory neurons. We showed 13 chromosomes which contain 25Kb bin resolution SnapFISH-identified loops. Note that imaging data in chr2, chr3, chr6, chr8 and chr15 contain missing data, which are showed in grey vertical and horizontal lines. In each heatmap, the dashed black cycle and the grey pixel in the lower left represents the location of the SnapFISH-identified loops.

**Figure S7.**
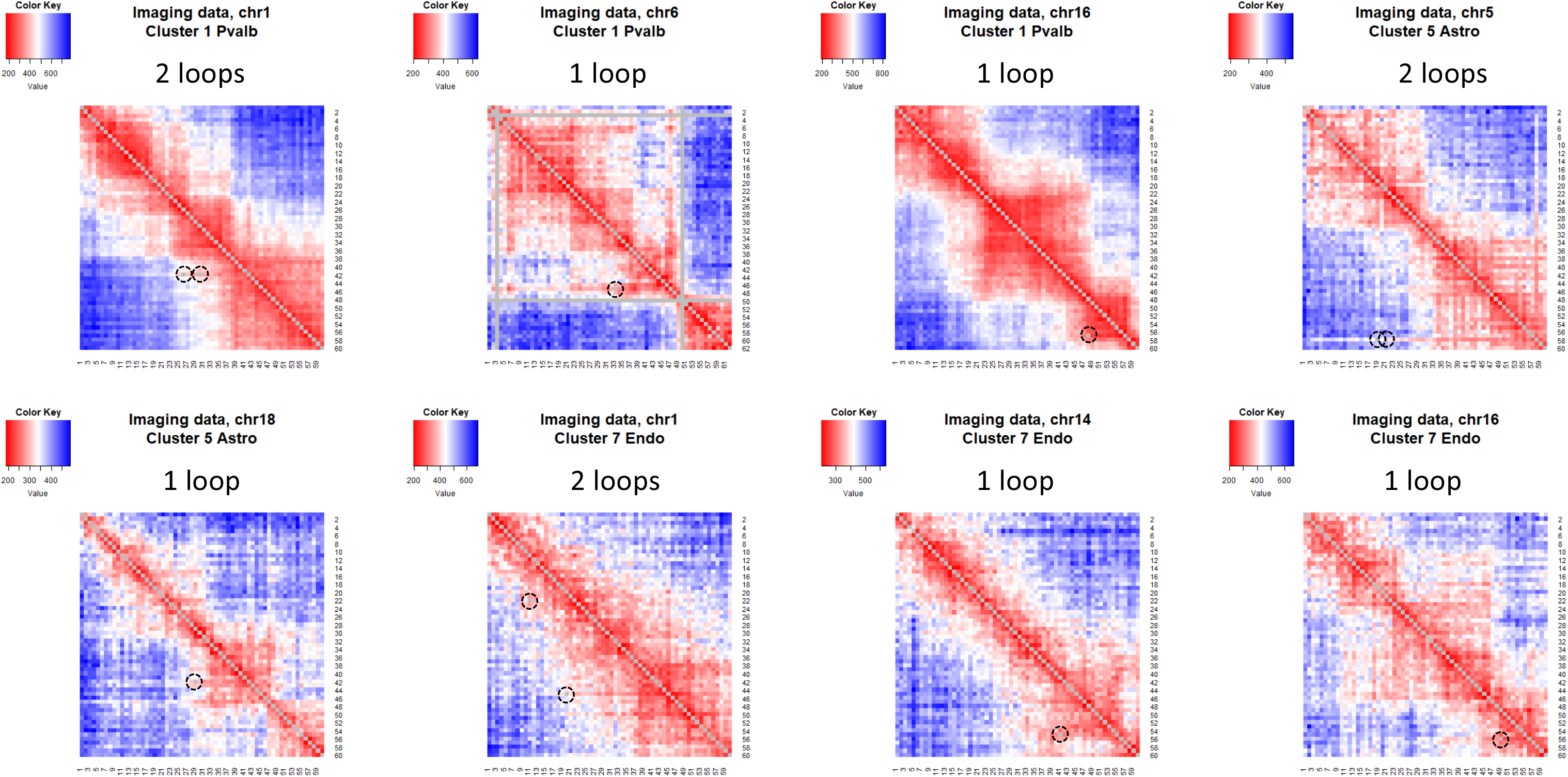
Heatmap of average Euclidean distance (unit: nm) from DNA seqFISH+ data in three additional mouse brain cell types (Pvalb, Astro and Endo). We showed chromosomes which contain 25Kb bin resolution SnapFISH-identified loops. Note that imaging data in chr6 contain missing data, which are showed in grey vertical and horizontal lines. In each heatmap, the dashed black cycle and the grey pixel in the lower left represents the location of the SnapFISH-identified loops.

**Figure S8.**
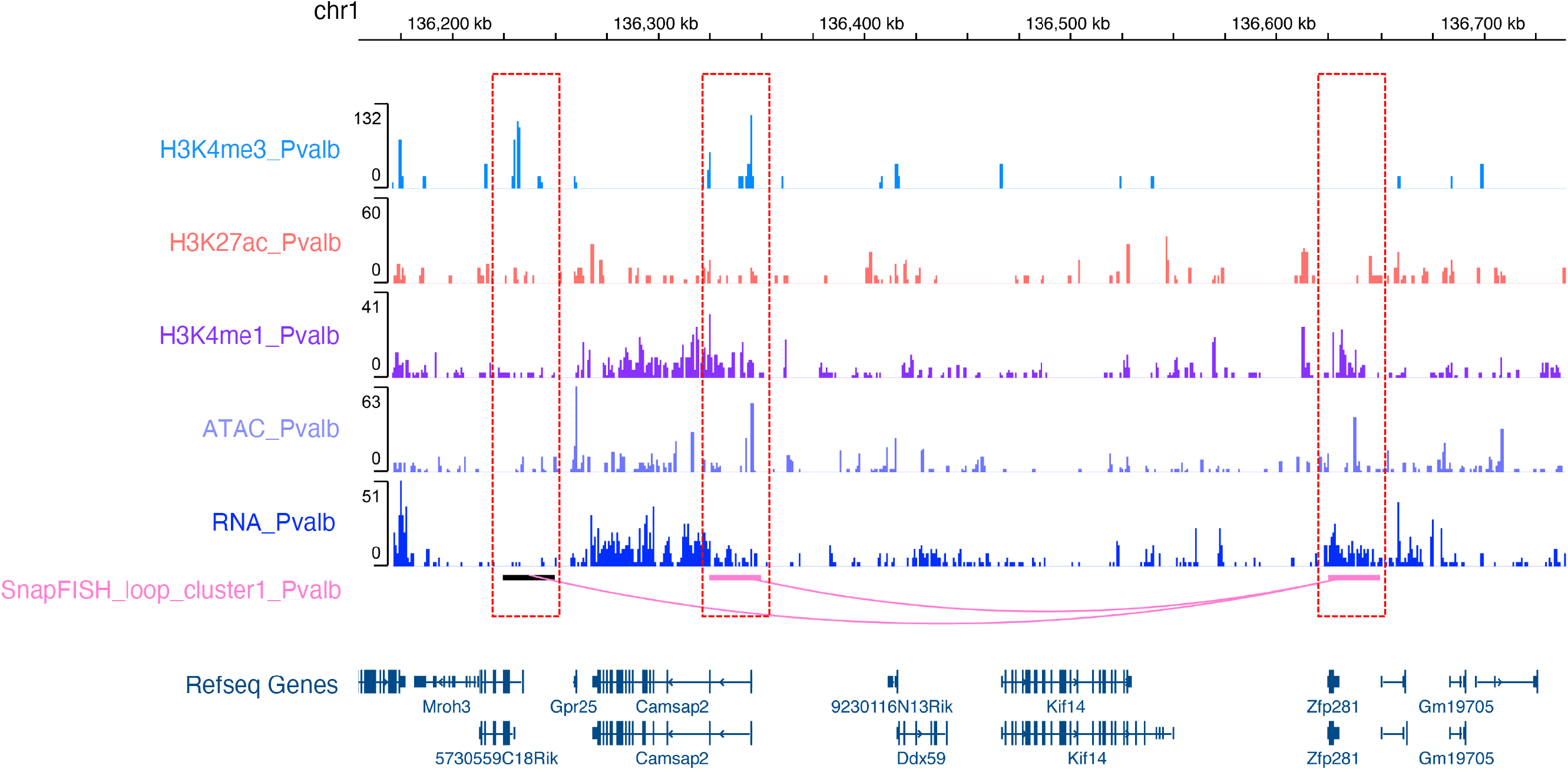
SnapFISH identified two enhancer-promoter loops in chr1 from DNA seqFISH+ data in mouse inhibitory neurons expressing parvalbumin (Pvalb). The top five tracks are H3K4me3, H3K27ac, H3K4me1, ATAC-seq and RNA-seq data from inhibitory neurons expressing parvalbumin collected from mouse frontal cortex tissue (Pvalb for short)^30^. The bottom two tracks are the SnapFISH-identified 25Kb bin resolution loop and the Refseq gene annotation. The left anchor (dashed red box on the left) and the middle anchor (dashed red box in the middle) are at the promoter of gene *Mroh3* and the promoter of gene *Camsap2*, respectively, and both contain strong H3K4me3 ChIP-seq peak. The right anchor (dashed box on the right) overlaps H3K4me1 and H3K27ac peaks, as well as ATAC-seq peaks.

**Figure S9.**
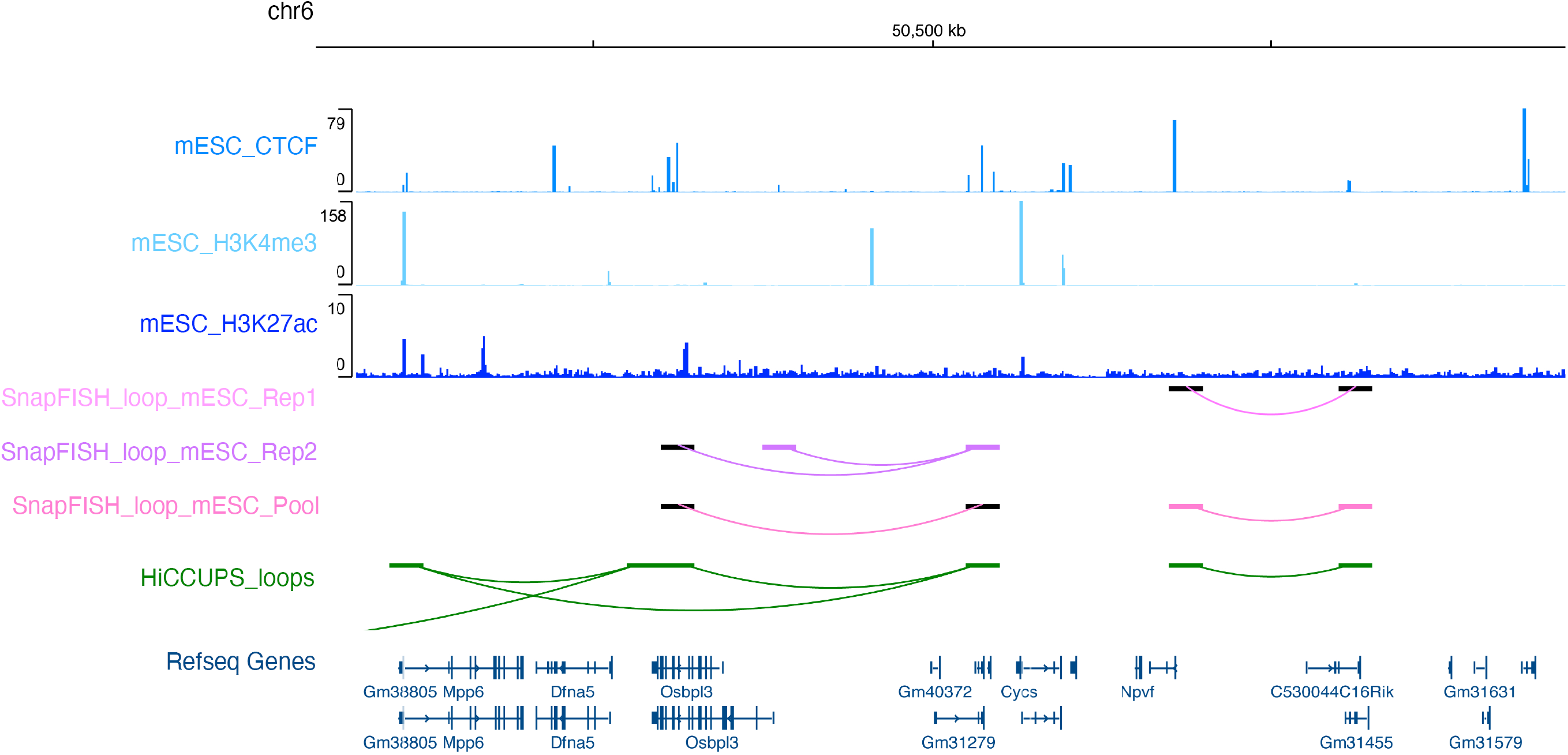
SnapFISH identified loops in chr6 from two biological replicates of mESC DNA seqFISH+ data. The top three tracks are mESC CTCF, H3K4me3 and H3K27ac ChIP-seq data. The middle four tracks are SnapFISH-identified loops from replicate #1 (Rep1), replicate #2 (Rep2), pooled data, and HiCCUPS-identified loops from mESC bulk Hi-C data. All loops are at 25Kb bin resolution. The bottom track is the Refseq gene annotation.

**Figure S10.**
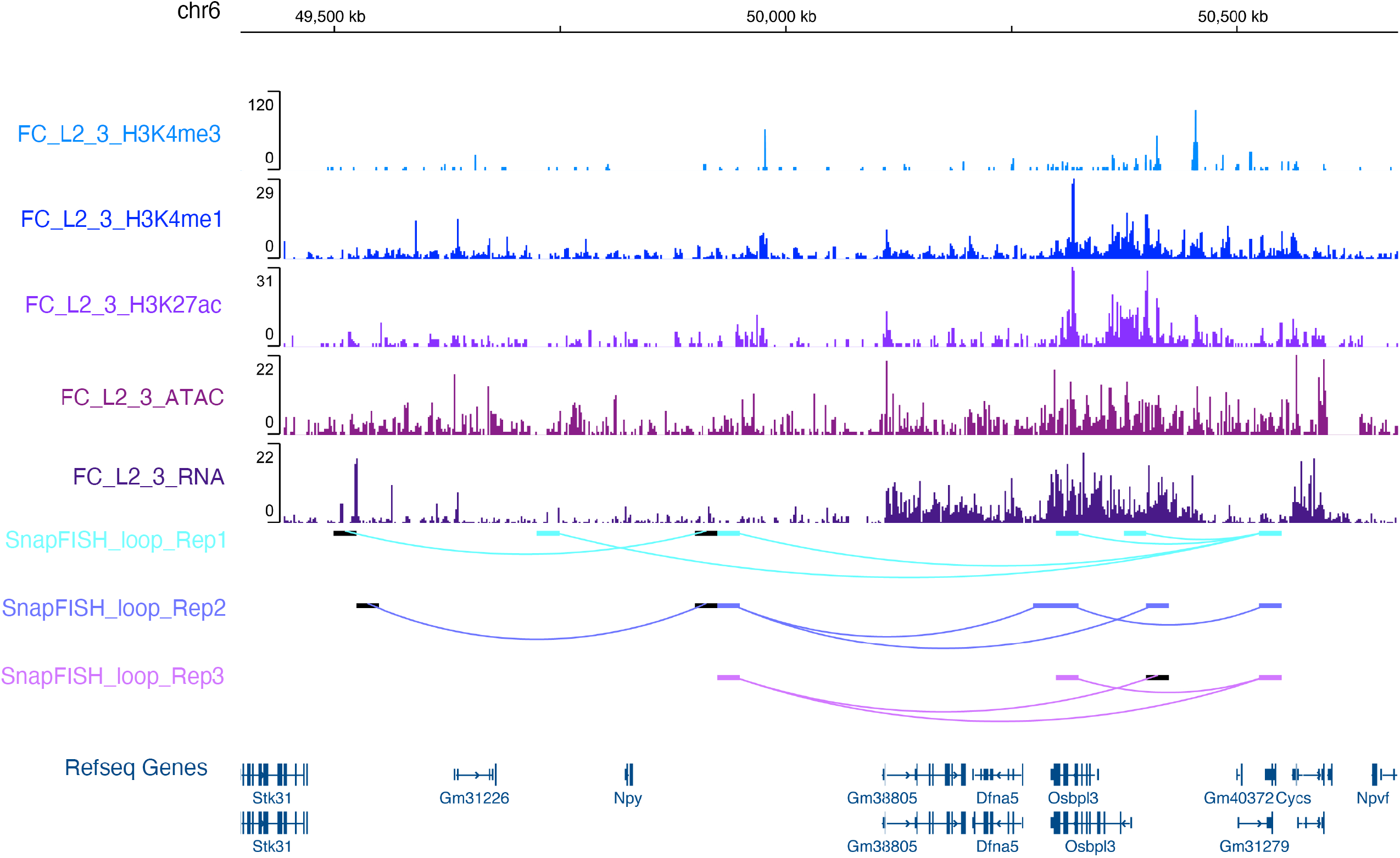
SnapFISH identified loops in chr6 from three biological replicates of DNA seqFISH+ data in mouse excitatory neurons. The top three tracks are SnapFISH-identified loops in replicate #1 (Rep1), replicate #2 (Rep2) and replicate #3 (Rep3), respectively. All loops are at 25Kb bin resolution. The bottom track is the Refseq gene annotation.

**Figure S11.**
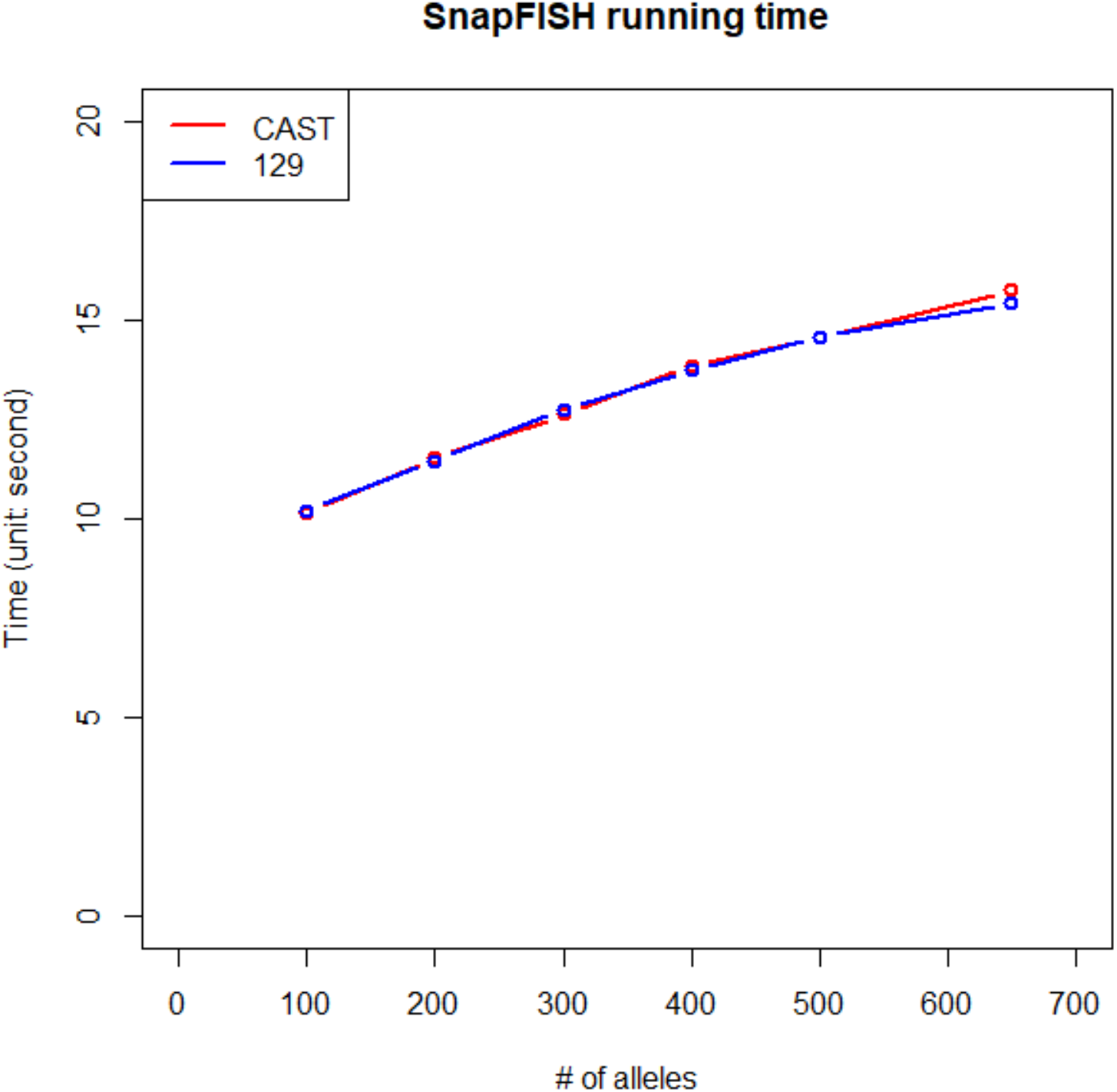
SnapFISH running time with different numbers of alleles.

## Table titles

**Table S1. Sensitivity of SnapFISH using different numbers of alleles**. Column K is the indicator of whether SnapFISH-identified loop is overlapped with the known *Sox2* enhancer-promoter loop. (A) the CAST allele. (B) the 129 allele.

**Table S2. Sensitivity of SnapFISH using 472 alleles with three different levels of targeted segment detection efficiency**. Column L is the indicator of whether SnapFISH-identified loop is overlapped with the known *Sox2* enhancer-promoter loop. **A.** the CAST allele. **B.** the 129 allele.

**Table S3. Chromatin regions with DNA seqFISH+ data for mESCs and mouse excitatory neurons**.

**Table S4. SnapFISH identified chromatin loops from DNA seqFISH+ data. A.** 6 loops in mESCs. **B.** 28 loops in mouse excitatory neurons. **C.** 4 loops in inhibitory neurons expressing parvalbumin (Pvalb for short), 3 loops in astrocytes (Astro for short) and 4 loops in endothelial cells (Endo for short).

**Table S5. Evaluation of the sensitivity of SnapFISH in mESC DNA seqFISH+ data, using 16 HiCCUPS-identified loops from mESC bulk Hi-C data as the reference loop list**.

**Table S6. SnapFISH-identified loops in the two biological replicates of mESC DNA seqFISH+ data**.

**Table S7. SnapFISH-identified loops in the three biological replicates of DNA seqFISH+ data in mouse excitatory neurons**.

## Methods

### Definition of local neighborhood regions

For both 5Kb bin resolution multiplexed DNA FISH data in mESCs, 5Kb bin resolution OCRA data in mESCs, and 25Kb bin resolution DNA seqFISH+ data in mESCs and mouse excitatory neurons, we define the local neighborhood of a given targeted segment pairs as all of the identified pairs that fall within a square with 25Kb ~ 50Kb in 1D genomic distance from the targeted segment pair of interest. Specifically, **Figure S1** shows the definition of local neighborhood regions, which is similar to the definition that has been used in the HiCCUPS algorithm^4^, and our recently developed SnapHiC algorithm^18^. Specifically, the yellow, orange and purple areas represent the lower left, horizontal and vertical regions, respectively. The union of yellow, orange, purple and blue areas consist of the “square” region that define the local neighborhood.

### Identification of loop candidates

For each pair of targeted segments, we calculate the average Euclidean distance *d* across all cells. We further calculate the mean Euclidean distance within the square, lower left, horizontal and vertical regions within each cell, and then average across all cells, termed as *d_S_*, *d_LL_*, *d_H_*, *d_V_*. We define a pair of targeted segments as a loop candidate if and only if *d_S_*/*d* > 1.1, *d_LL_*/*d* > 1.05, *d_H_*/*d* > 1.05, *d_V_*/*d* > 1.05, *T* < –4 and *FDR* < 10% (**Figure 1D** and **Figure S1**).

### Identification of loop summits

We group nearby loop candidates within a pre-specified gap into clusters, where the gap is twice the size of the bin resolution. In other words, we defined the gap to be 10Kb for 5Kb resolution multiplexed DNA FISH data and 5Kb resolution OCRA data. We defined the gap to be 50Kb for 25Kb resolution DNA seqFISH+ data. We first remove singletons, and for the remaining clusters consisting of more than two loop candidates, we define the ones with the most significant FDR as the loop summits. For 5Kb resolution multiplexed DNA FISH data and ORCA data, we use the loop summits as the final output. For 25Kb resolution DNA seqFISH+ data, we treat both loop summits and singletons as the final output.

### Overlap of SnapFISH-identified loop with *Sox2* enhancer-promoter loop

We defined chr3:34,645,000-34,655,000 and chr3:34,755,000-34,765,000 as the two 10Kb bins containing *Sox2* promoter and super-enhancer, respectively. We further defined a SnapFISH-identified 5Kb bin resolution loop as “overlapped” with the *Sox2* enhancer-promoter, if and only if the SnapFISH loop anchors are within 5Kb of *Sox2* promoter-super-enhancer loop anchors.

### Imaging data resource

1. **5Kb bin resolution multiplexed DNA FISH data from mESC at the *Sox2* gene locus.** We downloaded multiplexed DNA FISH data from 4D Nucleome data portal (https://data.4dnucleome.org/experiment-set-replicates/4DNESC5PKTQ9/), which are originally used in the Huang et al study^16^.
2. **5Kb bin resolution ORCA data from mESC at the *Sox2* locus.** ORCA data (Sox2_B1_T1_core.csv) are shared by Dr. Boettiger, which are originally used in the Mateo et al study^10^.
3. **25Kb bin resolution DNA seqFISH+ data from mESC.** We downloaded DNA seqFISH+ data from the website https://zenodo.org/record/3735329, and used two files (DNAseqFISH+25kbloci-E14-replicate1.csv and DNAseqFISH+25kbloci-E14-replicate2.csv). Such data are originally used in the Takei et al study^8^.
4. **25Kb bin resolution DNA seqFISH+ data from mouse cerebral cortex tissue.** We downloaded DNA seqFISH+ data from the website https://zenodo.org/record/4708112, and used the file (TableS8_brain_DNAseqFISH_25kb_voxel_coordinates_2762cells.csv). Such data are originally used in the Takei et al study^9^.

### mESC ChIP-seq data resource

mESC H3K4me3 ChIP-seq data is from our previous study^33^. mESC H3K27ac ChIP-seq data is downloaded from the ENCODE website: https://www.encodeproject.org/experiments/ENCSR000CGQ/. mESC CTCF ChIP-seq data is downloaded from the ENCODE website: https://www.encodeproject.org/experiments/ENCSR000CCB/.

## Notes

### Competing Interest Statement

Bing Ren is a cofounder and shareholder of Arima Genomics, Inc. and Epigenome Technologies, Inc. The remaining authors declare no competing interest.

