## Supplementary Materials for "SnapFISH: a computational pipeline to identify chromatin loops from multiplexed DNA FISH data"

**Supplementary Information**

**Section 1: Applying SnapFISH to DNA seqFISH+ data in three additional mouse brain cell types.**

Takei et al study^1^ contains DNA seqFISH+ data of 2,762 cells from nine major cell type clusters. We have showed that SnapFISH can identify enhancer-promoter loops in mouse excitatory neurons (**Figure 2C** and **Figure 2D**), which is the largest cell type cluster consisting of 1,895 cells (3,790 alleles). We then evaluated the performance of SnapFISH on other cell type clusters with less number of cells. Since our previous analysis has showed that SnapFISH can only identify the *Sox2* enhancer-promoter loop with more than 200 alleles (**Figure S3** and **Table S1**), we applied SnapFISH to DNA seqFISH+ data generated from three additional major cell type clusters with more than 200 alleles, including inhibitory neurons expressing parvalbumin (Pvalb for short, 155 cells, 310 alleles), astrocytes (Astro for short, 152 cells, 304 alleles) and endothelial cells (Endo for short, 240 cells, 480 alleles). All the remaining five cell type clusters consist of less than 200 alleles.

Notably, the targeted segment detection efficiency is 39.5%, 38.6% and 19.6% for Pvalb, Astro and Endo, respectively, which are comparable (Pvalb and Astro) or much lower (Endo) than the 43.1% targeted segment detection efficiency in excitatory neurons. The low targeted segment detection efficiency and the small number of alleles in these three cell types may results in the limited sensitivity of loop calling. As expected, SnapFISH identified 4, 3 and 4 loops in Pvalb, Astro and Endo, respectively (**Table S4C** and **Figure S7**), which are much less than 28 loops identified from excitatory neurons. We also used Paired-Tag data^2^ to annotate these SnapFISH-identified loops. **Figure S8** showed an illustrative example of two enhancer-promoter loops in chr1 in Pvalb, where the loop connects the putative enhancer near gene *Zfp281* to the promoter of gene *Mroh3* and the promoter of gene *Camsap2*. This example showcased that SnapFISH can identify enhancer-promoter loops in Pvalb with less number of cells compared to the largest cell type excitatory neurons.

**Section 2. Evaluation the reproducibility of SnapFISH.**

We first evaluated the reproducibility of SnapFISH using 25Kb bin resolution DNA seqFISH+ data generated from mESCs^3^, which consists of two biological replicates. Replicate #1 contains 201 cells (i.e., 402 alleles) with targeted segment detection efficiency 58.6%, while Replicate #2 contains 245 cells (i.e., 490 alleles) with targeted segment detection efficiency 70.6%. Applying SnapFISH to these two replicates separately identified 1 loop and 6 loops in Replicate #1 and Replicate #2, respectively (**Table S6**). The more loops detected from Replicate #2 might be due to the fact that Replicate #2 contains more cells with higher targeted segment detection efficiency. The only one loop in chr6 identified from Replicate #1 is not reproducible with two loops in chr6 identified from Replicate #2. We looked into these non-reproducible loops in chr6, and found that they are overlapped with different HiCCUPS loops from mESC bulk Hi-C data (**Figure S9**). Notably, SnapFISH loops in Replicate #2 contains one false positive which is not overlapped with HiCCUPS loops, while SnapFISH loops from the pooled data (Replicate #1 + Replicate #2) do not contain such false positive. We also checked the accuracy of 4 additional SnapFISH loops in Replicate #2, and found that they are all overlapped with HiCCUPS loops (**Table S6**). Therefore, the precision is 100% and 83.3% for SnapFISH loops in Replicate #1 and Replicate #2, respectively. Taken together, our data show that SnapFISH can identify loops with high accuracy in each biological replicate. The low loop reproducibility underscores the high cell-to-cell variability of chromatin loops among the cell population.

We further evaluated the reproducibility of SnapFISH using 25Kb bin resolution DNA seqFISH+ data generated from mouse excitatory neurons^1^, which consists of three biological replicates. Replicate #1 contains 1,077 alleles with targeted segment detection efficiency 41.7%, Replicate #2 contains 1,072 alleles with targeted segment detection efficiency 42.8%, while Replicate #3 contains 1,567 alleles with targeted segment detection efficiency 46.2%. Applying SnapFISH to these three replicates separately identified 18 loops, 36 loops and 26 loops in Replicate #1, Replicate #2 and Replicate #3, respectively (**Table S7**). For each replicate, we defined the reproducibility as the number of loops are overlapped (up to 25Kb gap) in at least one of two other replicates. Based on such definition, 11 out 18 loops in Replicate #1 (61.1%), 15 out of 36 loops in Replicate #2 (41.7%) and 16 out of 26 loops in Replicate #3 (61.5%) are reproducible. **Figure S10** shows an illustrative example in chr6, where 3 out 5 loops in Replicate #1 (60.0%), 3 out of 4 loops in Replicate #2 (75.0%) and 3 out of 3 loops in Replicate #3 (100.0%) are reproducible. Collectively, our data showed that SnapFISH can identify reproducible loops among three biological replicates. The loops specific to only one replicate may be due to the high cell-to-cell variability of chromatin loops among the cell population, as what we observed in mESC DNA seqFISH+ data.

**Section 3. Time and memory cost of SnapFISH.**

We evaluated the time and memory cost of SnapFISH, using 5Kb bin resolution mESC multiplexed DNA FISH data with different numbers of the CAST alleles and the 129 alleles (i.e., # of cells = 100, 200, 300, 400, 500, and 649). **Figure S11** showed that SnapFISH is computationally efficient, which only takes ~15 seconds to analyze 649 alleles, and the running time increases approximately linearly with the number of alleles. In addition, SnapFISH requires ~90Mb memory for different numbers of CAST alleles and 129 alleles. All computation tasks are performed in the single compute node with a 2.40 GHz Intel processor and 256 Gb RAM.
